# Structural Dynamics of RNA Polymerase II During Nucleotide Addition Cycle

**DOI:** 10.64898/2026.06.04.730248

**Authors:** Gangshun Yi, Qingrong Li, Hannah Holmberg, Shisheng Li, Daniel K. Clare, Dong Wang, Peijun Zhang

**Author notes:** These authors contributed equally (Gangshun Yi, Qingrong Li).

## Abstract

RNA polymerase II (RNAPII) drives gene expression through iterative nucleotide addition cycles (NACs) comprising translocation, substrate binding, and catalysis. The lack of pre-catalysis and post-catalysis intermediates has precluded a complete mechanistic understanding of the NAC. Here we present 43 cryo-EM structures capturing distinct stages of the *S. cerevisiae* RNAPII elongation complex (EC) NAC, including previously intractable transition intermediates. We establish a continuous spectrum of RNAPII EC structural dynamics during the NAC, which can be divided into two coordinated phases: a substrate-induced EC tightening phase and a post-catalysis EC relaxation phase. For the substrate-induced EC tightening phase, the substrate binding initiates allosteric conformational changes across the entire RNAPII EC, including TL folding, funnel closure, clamp closure, transcription bubble ordering, and precise alignment of the RNA 3′-end with substrate to form a catalysis-competent configuration. For the post-catalysis EC relaxation phase, we captured the long-sought, short-lived post-catalysis product state and identified a series of intermediates that reveal a reverse conformational transition that facilitates rapid translocation. Together, our findings define a comprehensive structural and dynamic framework for RNAPII NAC, yielding a “molecular movie” of RNAPII in action and revealing a fundamental principle by which the enzyme balances speed and fidelity through coordinated conformational dynamics.

## Introduction

RNA polymerase II (RNAPII) is a central enzyme for gene transcription. During transcription, RNAPII selects the correct substrate for reaction and undergoes significant conformational changes in an iterative nucleotide addition cycle (NAC)^1–7^. NAC can be divided into three major stages: translocation, substrate binding, and catalysis (Fig. 1a–b)^3,4^. During translocation^8^, RNAPII moves one base pair forward from the pre-translocation state^9^ to the post-translocation state^10,11^, in which the template base is loaded at i+1 position and the active site is available for substrate binding^11,12^. During substrate binding, RNAPII selects and binds the cognate substrate at the addition site (A-site) to form a Watson-Crick base pair with template base at i+1 position^12,13^. Previous research has shown that the trigger loop (TL) (Rpb1 residue Met1063–Val1107), a conserved structural motif, switches from an open to a closed state upon correct substrate binding, which is essential for RNAPII substrate selection and subsequent catalysis^12–15^. During catalysis, a phosphodiester bond forms between the incoming substrate and the 3′-end of RNA. Following bond formation, the TL is thought to transition from a closed to an open state, releasing byproduct pyrophosphate from the active site and returning the complex to the pre-translocation state, ready for the next cycle.

**Figure 1.**
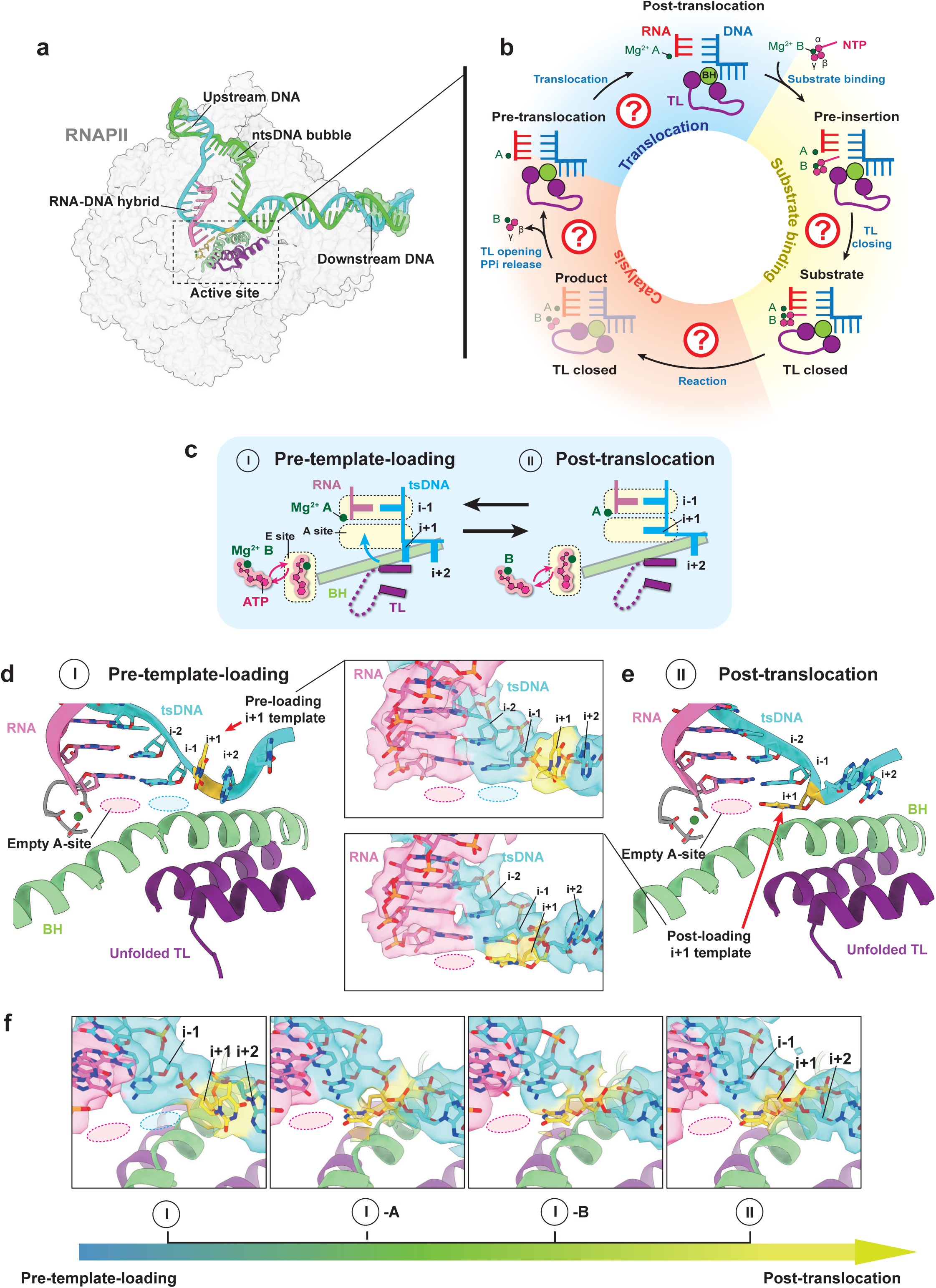
Novel RNAPII EC structure at the pre-template-loading state and conformational transitions. (a) Representative structure of the RNAPII EC. The structure is shown from Dataset 2 (frame 20, state 5) and is presented as a representative conformation of the RNAPII EC. RNAPII is colored in silver, with tsDNA (cyan), ntsDNA (lime), and RNA (hot pink). The active center is highlighted by a dashed box. Key active center elements, including the BH (light green) and TL (purple), are shown. The substrate ATP is depicted in gold. Abbreviations, residue ranges, and color codes for RNAPII EC elements are summarized in Table S6. (b) Schematic illustrating sequential conformational changes within the active center of RNAPII during the nucleotide addition cycle. Four of the five states have been resolved previously by cryo-EM or X-ray crystallography (solid colors). The previously unresolved product state is depicted as a transparent cartoon. The nucleotide addition cycle is categorized into three sections highlighted by distinct background colors: translocation (blue), substrate binding (yellow green), and catalysis (orange). Unknow mechanisms of conformational transition are indicated as question marks. Domain-specific coloring is consistent with panel a. (c) Cartoon representation detailing the newly identified pre-template-loading state (state I) and its transition to the post-translocation state (state II), observed in both the RNAPII EC and RNAPII EC substrate binding datasets. (d–e) Cryo-EM structures of RNAPII EC in pre-template-loading state (state I, panel d) and post-translocation state (state II, panel e) derived from the RNAPII EC substrate-free. Insets highlight cryo-EM density corresponding to the loading of the i+1 template nucleotide (contoured at 3-σ). Coloring of RNAPII EC elements matches the scheme depicted in panel c. (f) Cryo-EM maps displaying dynamic density changes during the translocation of the i+1 template nucleotide (contoured at 3-σ).

Despite significant progress over the past 25 years^2,5–7^, our understanding of the structural dynamics of the NAC remains limited, relying largely on four static snapshots of the translocation and substrate-binding steps (Fig. 1b)^3,4^. The critical process of catalysis, one of the central stages of the NAC, is not understood due to the absence of a high-resolution product structure upon catalysis (Fig. 1b, shaded orange). This product state is short-lived and very challenging to capture using canonical structural approaches. Furthermore, the molecular basis of dynamic transitions between these states of the NAC remains elusive. Long-range allosteric conformational changes triggered by substrate binding and catalysis, beyond the TL opening and closing, are poorly understood. Several fundamental questions remain unanswered: Is template loading synchronized with upstream DNA/RNA movement during translocation? How does TL get folded to a closed conformation upon substrate binding and then unfolded after catalysis? What is the temporal relationship between pyrophosphate release and TL opening? Whether and how does substrate binding lead to allosteric conformational changes in structural motifs beyond the TL, and how are these motions communicated?

Here we report cryo-EM structures capturing distinct stages of the *S. cerevisiae* RNAPII NAC, including previously intractable intermediates, and reveal dynamic conformational transitions across all stages of the cycle. In translocation, we identified a previously uncharacterized pre-template-loading state. In substrate binding, we resolved multiple novel intermediates that trace a near-continuous trajectory of TL folding, from fully open to partially folded to fully folded conformations. The substrate binding initiates long-range allosteric conformational changes across the entire RNAPII elongation complex (EC), which we refer to as “substrate-induced EC tightening.” These include TL folding, funnel closure, clamp closure, transcription bubble ordering, and precise alignment of the RNA 3′-end with the incoming substrate to form a catalysis-ready configuration. In catalysis, we captured the long-sought, short-lived post-catalysis product state and identified a series of intermediates that reveal a reverse conformational transition, which we termed “post-catalysis EC relaxation,” as RNAPII resets to a relaxed, translocation-ready state. Taken together, our results establish a coherent and comprehensive structural dynamic framework for the complete NAC, yielding a “molecular movie” of RNA polymerase II in action and revealing mechanistic insights that resolve long-standing questions in transcription.

## Results

To achieve a comprehensive understanding of the RNAPII EC NAC, we designed four cryo-EM samples to dissect the key steps of the *S. cerevisiae* RNAPII elongation cycle: a substrate-free RNAPII EC containing a 3′-deoxy (3′-dOH) 9-mer RNA to capture the different translocation states (Dataset 1) (Extended Data Fig. 1, Table S1); a RNAPII EC containing a 3′-dOH 9-mer RNA and ATP substrate to probe substrate binding states (Dataset 2) (Extended Data Fig. 2, Table S2–S3); a RNAPII EC with a regular 9-mer RNA primer and NTP substrates to enable completion of full NAC cycles (Dataset 3) (Extended Data Fig. 3–4, Table S4); and a RNAPII EC containing a long 3′-dOH RNA (19-mer) and ATP substrate to mimic a mature EC (Dataset 4) (Extended Data Fig. 5, Table S5). We employed monodispersed single particle streptavidin (mspSA) affinity grids for cryo-EM sample preparation, which effectively mitigated air-water interface effects and reduced RNAPII orientation bias, thereby markedly enhancing resolution^16^. In both Datasets 2 and 4, binding of ATP primes the EC complex for catalysis, but the absence of a 3′ hydroxyl group prevents nucleotide addition. For Dataset 3, to sustain continuous NAC turnover and capture transient intermediates directly on cryo-EM grids, we employed a substrate-gradient affinity grid (SGAG) strategy (see Methods and Extended Data Figs. 3 and 4 for details). From these datasets, we reconstructed 43 cryo-EM maps of RNAPII EC intermediates at resolutions ranging from 2.5 Å to 4.3 Å and built 31 corresponding atomic models (Tables S1–S5). Abbreviations, residue ranges, and color codes for RNAPII EC elements used in this study are listed in Table S6. Collectively, these structures provide a comprehensive view of RNAPII structural dynamics, revealing conformational changes triggered by substrate binding and catalysis during the NAC (Summarized in Movie S1).

### New template loading intermediates reveal unsynchronized RNAPII translocation

Previous studies have shown that multi-subunit RNAPs (including RNAPII) can spontaneously oscillate among translocation states in the absence of NTPs *in vitro*^17–24^. In addition to the previously characterized post-translocation state^10^, we captured several new translocation intermediates in the substrate-free dataset (Dataset 1). The first intermediate, termed the pre-template-loading state (State I), features an upstream-shifted RNA-DNA hybrid, while the i+1 template base has not yet crossed the bridge helix (Rpb1 residues Pro810–Ile848) or fully loaded into the active site (Fig. 1c–d). The position of the i+1 base in this pre-template-loading state differs markedly from its canonical location in the post-translocation state^10^ (State II, Fig. 1b–d). We also identified two additional intermediates, States I-A and I-B, which capture progressive density shifts of the i+1 nucleotide as it transitions from State I to the post-translocation conformation (State II) (Fig. 1e).

Intriguingly, the pre-template-loading state was also observed in the RNAPII EC-substrate-binding dataset (Dataset 2; Fig. 2a–b; Movie S2), indicating that the transition from pre-template-loading to post-translocation states can occur with or without a substrate in the entry site (E-site). Electron density corresponding to the ATP triphosphate moiety was detected at the E-site, whereas the nucleobase was unresolved, likely due to conformational flexibility. These observations suggest that ATP can associate with the E-site prior to template base loading.

**Figure 2.**
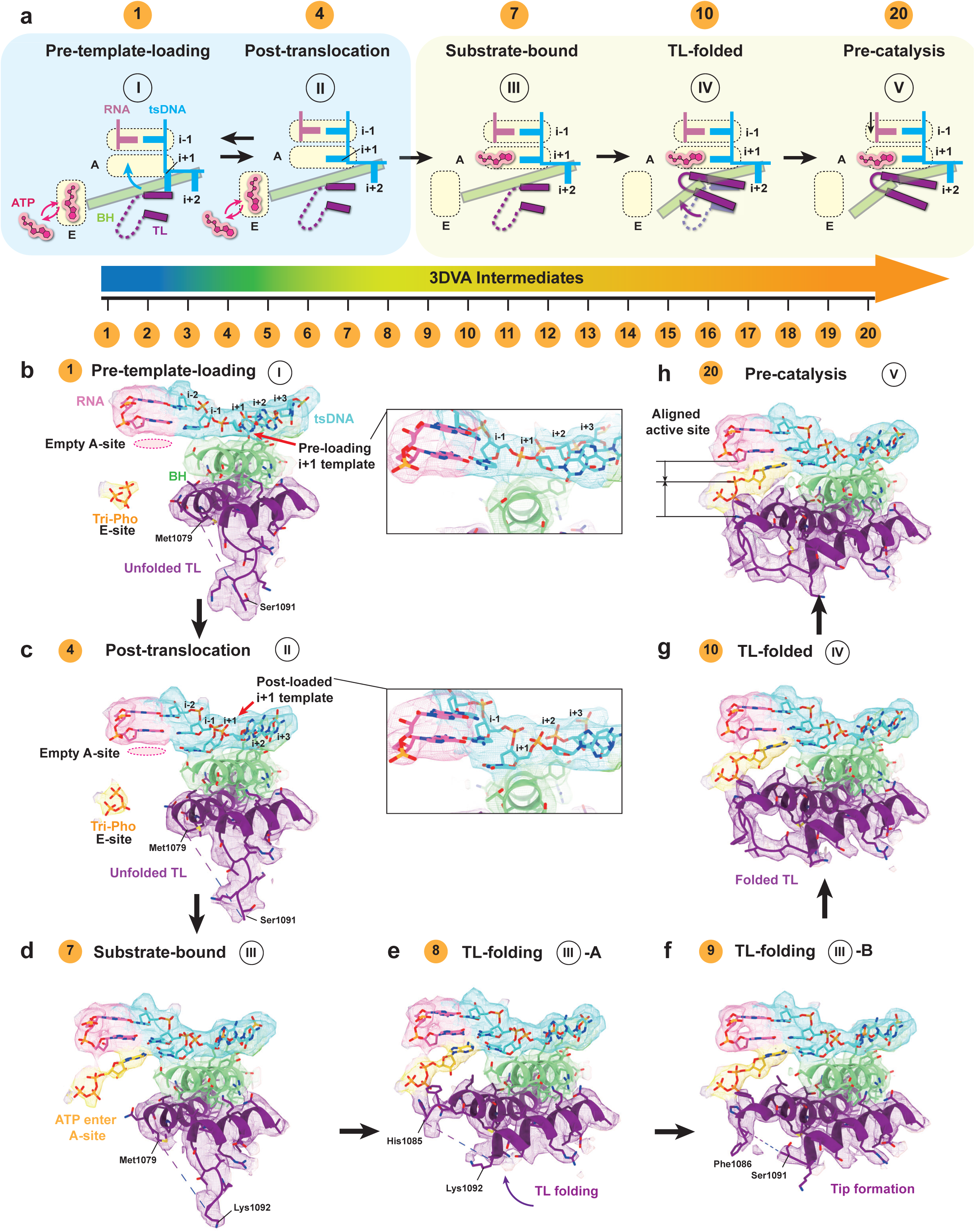
Structural dynamics of the TL folding and substrate loading. (a) Cartoon scheme illustrating the conformational dynamics and state transition in RNAPII’s active site during the substrate loading. A dynamic spectrum containing 20 intermediates (PC0) are shown in the lower panel. Cartoons schemes for five key states of transcription are shown in upper panel, including the pre-template-loading (State I, Frame 1), post-translocation (State II, Frame 4), substrate-bound (State III, Frame 7), TL-folded (State IV, Frame 10), and pre-catalysis (State V, Frame 20) states. RNAPII domains involved conformational dynamics are shown, including the TL (purple), bridge helix (light green), RNA primer & ATP substrate (hot pink), and template DNA (cyan). Frame indexes of key states are indicated and corresponding to the dynamic spectrum. (b-h) Cryo-EM structures of transcriptional states corresponding to the dynamic spectrum in panel (a), including five key states (b, c, d, g and h) as upper panel (a) shown and two intermediate states demonstrating the stepwise folding of TL (e and f). The color codes for RNAPII domains are correspond to the cartoon scheme in panel (a). The cryo-EM densities are shown at a uniform contour level of 0.12 (4-σ). Detailed structures of the i+1 template nucleotide in the pre-template-loading (b) and post-translocation (c) states, highlighting differences in the nucleotide positioning between these two states.

In the same dataset, we also captured the post-translocation state (State II), in which ATP remains bound at the E-site and has not yet transferred into the A-site (Fig. 2c). We further identified a third intermediate (State III), the substrate-bound state (Fig. 2d), where RNAPII adopts a post-translocation conformation, ATP occupies the A-site, and the TL remains open. Together, these structures (States I–III) suggest that the substrate initially binds non-specifically at the E-site, and that correct base pairing occurs only after template loading, guiding the selective transfer of the nucleotide into the A-site.

### Conformational dynamic intermediates upon substrate binding

To capture substrate-bound RNAPII intermediates, we utilized an RNA scaffold containing a 3′-dOH modification in Dataset 2, allows nucleotide binding but prevents catalysis. Conformational changes triggered by substrate binding were analyzed using three-dimensional variability analysis (3DVA)^25^, which applies a linear subspace model to cluster and sequentially order particle conformations along principal components (PCs) (Table S2), and complementary 3D classification (Table S3). The convergence of these independent analyses (Extended Data Fig. 6b–d; Tables S2, S3) reinforces the reliability of the captured conformational intermediates. The primary principal component (PC0) defined the dominant trajectory induced by substrate binding, referred to here as the “substrate-induced EC tightening” (Figs. 2–5). This trajectory reveals a coordinated structural transition, including TL folding (Movie S3), funnel closure (Movie S4), clamp closure (Movie S5), transcription-bubble stabilization (Movie S5), and active site alignment (Movie S6).

**Figure 3.**
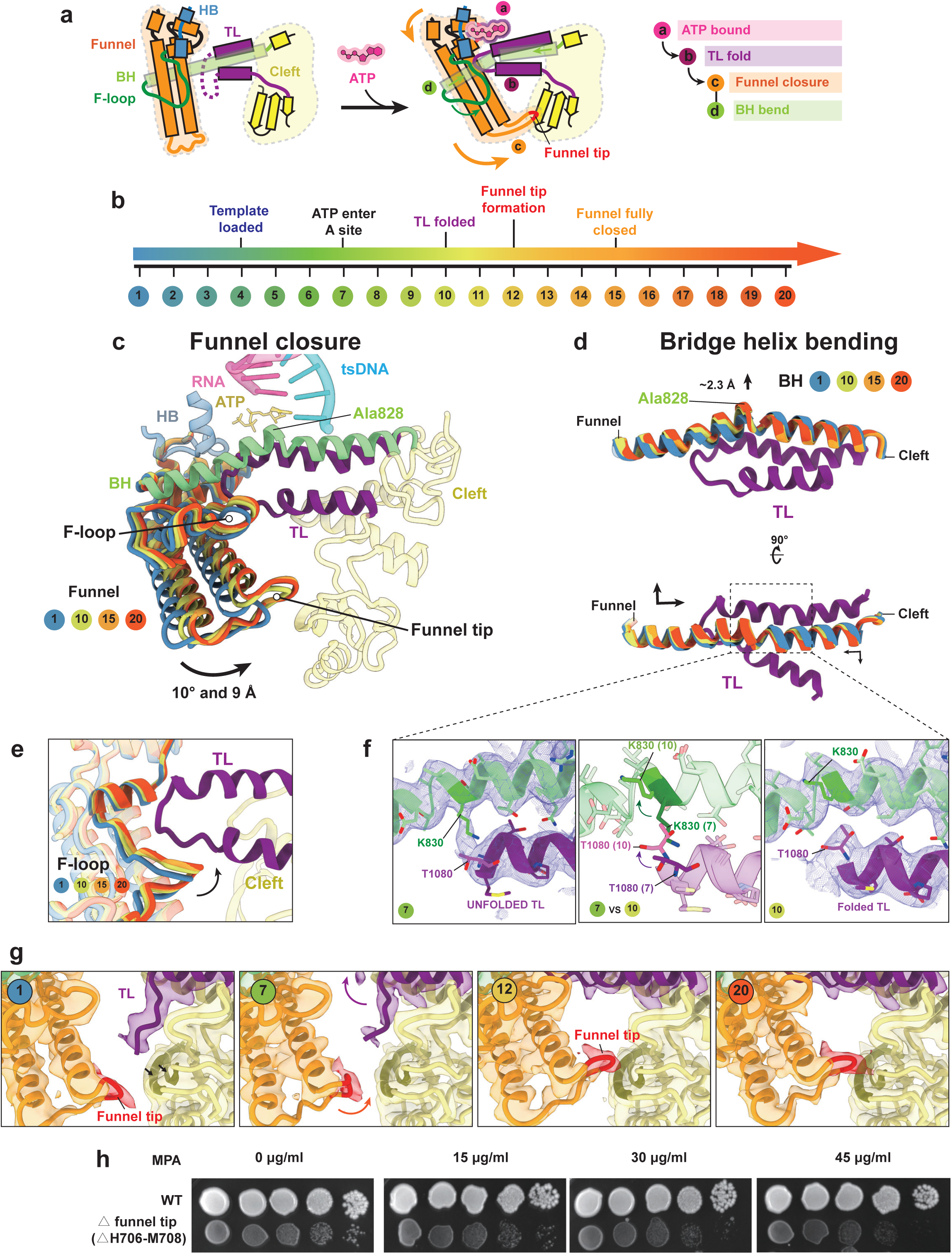
TL folding coordinates with BH bending and funnel closure. (a) Schematic illustration depicting the stepwise conformational changes associated with substate ATP binding, TL folding, BH bending and funnel closure. Color codes for different RNAPII domains are follows: TL (purple), BH (light green), HB (steel blue), funnel (orange), and cleft (yellow). The tip region of funnel domain is highlighted in red. (b) Frame spectrum represents dynamics across 20 cryo-EM intermediates (PC0) in Dataset 2 with a gradient color from blue (early) to red (late). Annotated Landmark events indicate key structural rearrangements along the trajectory. (c–e) Structural comparisons of the funnel domain (c), BH (d), and F-loop (e) across multiple 3DVA intermediates illustrate their coordinated conformational dynamics. Selected intermediates include frames 1, 10, 15, and 20. Arrows indicate the direction of domain movements along the trajectory from frame 1 to frame 20. Colors of the compared domains follow the 3DVA conformational spectrum defined in panel (b), while all other RNAPII elements are shown from frame 20 and colored as in panel (a). (f) Coupling dynamics between TL (purple) folding and BH (light green) bending. Densities and stick models of the unfolded (left, Frame 7) and folded (right, Frame 10) TL correspond to region marked in panel (d). Cryo-EM densities are shown as blue mesh and contoured at 4-σ. Structural comparisons between the unfolded (transparent) and folded (solid) TL alongside the BH in the middle panel, emphasizing the rotational changes. (g) Structural dynamics of funnel tip formation and its insertion into cleft domain. Four cryo-EM intermediate structures (Frame 1, 7, 12 and 20) are displayed side-by-side, focusing on the funnel tip region. Cryo-EM structures are shown as ribbon models with associated cryo-EM electron densities (contoured at 4-σ). Black arrows in left panel indicates the binding region for funnel tip on cleft domain. Two related β-strands (Val1282–Val1291, Tyr1298–Asp1309) in the binding site are highlighted in olive. (h) Deletion of the Rpb1 funnel tip (H706–M708) impairs cell growth and confers sensitivity to MPA. Saturated cultures of wild-type and Rpb1 funnel tip-deleted cells were serially diluted 10-fold and spotted onto YPD plates containing the indicated concentrations of mycophenolic acid (MPA). Plates were incubated at 30°C for 4 days before imaging.

**Figure 4.**
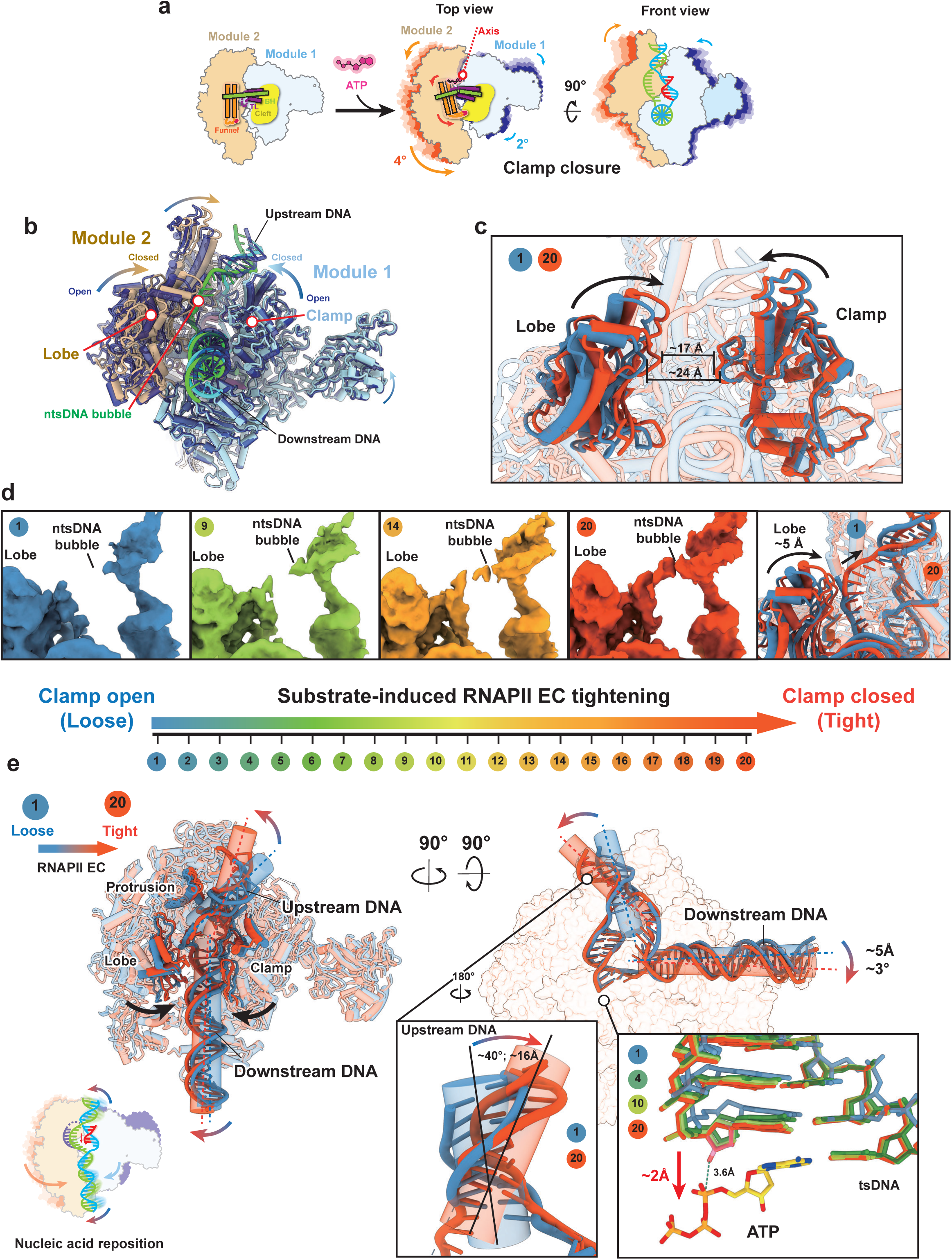
Substrate-induced clamp closure and nucleic acid scaffold repositioning. (a) Cartoon scheme of the substrate-induced clamp closure. Two major structural modules, module 1 (light blue) and module 2 (tan), exhibit opposing motions from pre-template load (State I, Frame 1) to pre-catalysis state (State V, Frame 20). The motion directions of clamp and nucleic acids are indicated by arrows. The rotation axis is indicated as the red dashed line. The core region in RNAPII EC plays an important role in this motion network, which consists of the TL (purple), BH (light green), funnel (orange) and cleft (yellow). (b) Structural comparison of RNAPII EC between state I and state V, illustrating global clamp closure. State I is shown in transparent blue, while state V is displayed in solid colors with domain-specific coloring consistent with panel (a). Arrows indicate the directions of module 1 and module 2 rotations. (c–e) Structural comparisons illustrating clamp closure (c), ntsDNA bubble stabilization (d), and nucleic acid scaffold repositioning (e) across multiple intermediate states in Dataset 2. Individual states used in each panel are indicated in the figures. Color coding of intermediate states follows the 3DVA frame spectrum shown in panel (d). Black arrows indicate the direction of module 1 and module 2 motions associated with clamp closure. In panel c, representative comparison of clamp closure using the lobe domain (module 2) and clamp domain (module 1) as markers. The closest distance between these domains is measured as indicated. In panel d, Cryo-EM density of the ntsDNA bubble region (contour level at 3-σ), showing progressive ordering and stabilization during the transition. In panel e, nucleic acid scaffold repositioning during RNAPII EC tightening. Selected elements lining the nucleic acid tunnel, including the protrusion, lobe, and clamp domains, are highlighted to illustrate tunnel narrowing. Upstream and downstream DNA trajectories are represented by shaded cylinders, with gradient-colored arrows (blue to red) indicating the direction of movement from the loose to tight state. Close-up views highlight upstream DNA repositioning and alignment of RNA primer and substrate ATP. In this view, a 3′-OH group was modeled onto the RNA primer; the distance between the RNA 3′-OH and the ATP α-phosphate is 3.6 Å. A red arrow indicates a ∼2 Å displacement of the RNA 3′-end during the RNAPII tightening. In the left corner of panel e, the cartoon schematic summarizes nucleic acid scaffold repositioning.

**Figure 5.**
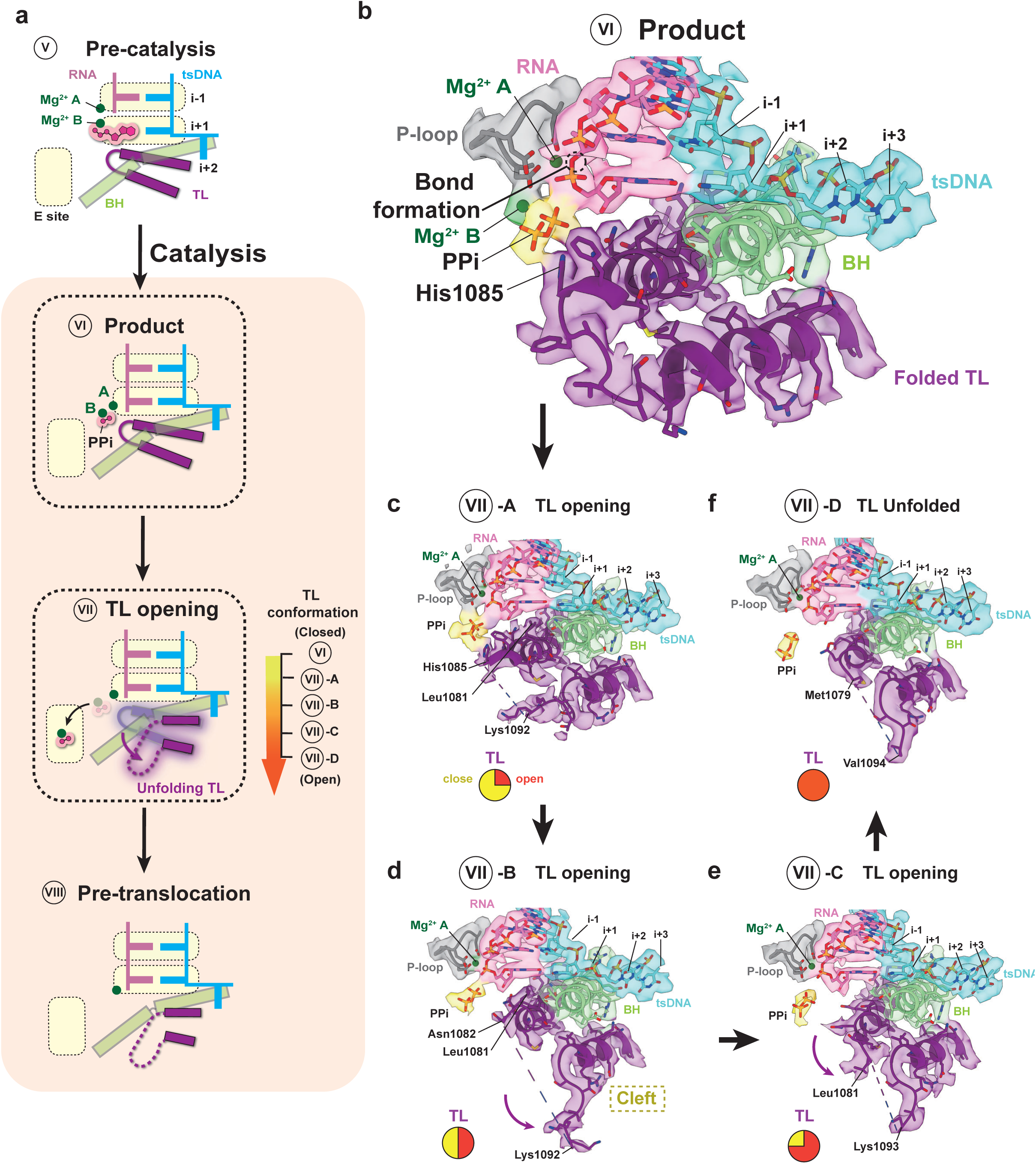
Cryo-EM structures of Pol EC at product state and TL-opening state after nucleotide addition. (a) Cartoon schemes illustrate the sequential transition of active center conformational states, including the pre-catalysis state (V), products state (VI), TL-opening state (VII), and pre-translocation state (VIII). A gradient spectrum adjacent to state VII depicts the TL conformational changes across four intermediate substates (VII-A, VII-B, VII-C, and VII-D), ranging from closed (yellow) to fully open (red). Color coding for distinct RNA polymerase II (RNAPII) domains is as follows: TL (purple), BH (light green), tsDNA (cyan), and RNA primer (hot pink). Arrows indicate the directions of TL movement and pyrophosphate (PPi) diffusion. (b–f) Cryo-EM structures of RNAPII EC captured at five distinct states: product state (VI, panel b) and TL-opening intermediate states VII-A (panel c), VII-B (panel d), VII-C (panel e), and VII-D (panel f). Cryo-EM density maps are contoured at 5-σ. The coloring and domain annotations correspond to those described in panel (a). Pie charts at the lower-left corners of panels (c–f) indicate the TL conformational distributions, with red representing open and yellow representing closed conformations.

Using particle subsets contributing to each frame, we reconstructed 20 cryo-EM structures representing distinct stages along this conformational trajectory, with global resolutions ranging from 3.3 Å to 4.3 Å (Fig 2, Extended Data Fig. 2 and Table S2). From these, we identified five primary conformational states: the pre-template-loading state (State I, Frame 1, Fig. 2b), post-translocation state (State II, Frame 4, Fig. 2c), substrate-bound state (State III, Frame 7, Fig. 2d), TL-folded state (State IV, Frame 10, Fig. 2g), and pre-catalysis state (State V, Frame 20, Fig. 2h). The strongest densities for both substrate ATP and the folded TL were observed in State V, indicating a stabilized, catalytically primed conformation. In addition, we resolved two previously uncharacterized TL sub-states (States III-A and III-B, Fig. 2e–f), providing further insight into the dynamic process of TL folding upon substrate engagement. The following sections describe the coordinated local and global conformational dynamics accompanying substrate binding.

### TL folding intermediates upon substrate engagement

A complete spectrum of TL folding intermediates induced by substrate binding was resolved through 3DVA (Movie S3). The transition of substrate ATP from the E-site to the A-site is tightly coupled with progressive TL folding. Prior to ATP loading into the A-site (Frames 1 to 6), the TL forms two helical segments (Met1063–Met1079 and Ser1091–Val1107), while the intervening region (Thr1080–Lys1092) remains disordered and lacks detectable density (Fig. 2b–c). Upon ATP engagement at the A-site (Frame 7, State III), the Met1063-Met1079 region remains flexible (Fig. 2d), consistent with the substrate-bound crystal structure (PDB: 2E2I)^12^.

Two intermediate states capture the stepwise process of TL folding (States III-A and III-B; Fig. 2e–f). In State III-A (Frame 8), the Lys1092 to Gly1097 segment detaches from the cleft and adopts a helical structure (Fig. 2e). In State III-B (Frame 9), the Met1079–His1085 region near the active center folds into a helix (Fig. 2f). Ordering of the loop between His1085 and Ala1090 forms a distinct tip, completing the fully folded TL conformation (Frame 10, State IV; Fig. 2g).

Following TL folding, the EC undergoes further rearrangements: The clamp continues to close, accompanied by precise repositioning of the nucleic acid scaffold and ATP. Densities for both ATP and the folded TL become stronger, reflecting a stabilized, catalytically poised conformation (Frame 20, State V, Fig. 2h). This ordered configuration is essential for efficient catalysis and ensures seamless progression through the transcription cycle.

### TL folding couples with BH bending and funnel closure

Substrate-induced TL folding results in a conformational transition that propagate through the RNAPII EC (Fig. 3a–b; Movies S4, S5). Upon folding, the TL tip inserts into a pocket formed by the hybrid-binding domain (HB), bridge helix (BH), and funnel domain (Rpb1 residue Ser663–Leu808 aa) (Fig. 3c), triggering a ∼10° rotation and ∼9 Å displacement of the funnel domain toward the cleft domain (Rpb1 residue Pro810–Asp871, His1059–His1140, Gly1275–Thr1394) (Fig. 3a–c). From Frame 1 (steel blue) to Frame 15 (orange), the funnel domain establishes new contacts with the folded TL tip, particularly via residues Ser754-Arg806 (Fig. 3c, 3e). This region, analogous to the F-loop in *E. coli* RNAP, is known to rearrange during TL-folding^26^. Beyond Frame 15, funnel closure reaches a largely converged state, with no substantial differences observed up to Frame 20 (Fig. 3c, 3e).

Before TL folding (Frame 1–7), the unfolded TL (Lys1092–Ser1096) blocks formation of the funnel tip and its interaction with the β-strand region of the cleft domain (Rpb1 residue Val1282–Val1291 and Tyr1298–Asp1309) (Fig. 3g). As folding progresses, the TL disengages, allowing the funnel tip (Rpb1 residue His706–Met708) to insert and form a new stabilizing interface after Frame 12 (Fig. 3c, 3g). This funnel tip appears to act as a conformational “lock” that reinforces the TL-closed, catalytically competent state. Notably, Gly707 within this tip is highly conserved (Extended Data Fig. 7). Deletion of the funnel tip impairs yeast growth, evidenced by smaller colonies (Fig. 3h), and increases sensitivity to mycophenolic acid (MPA), a nucleotide-depleting agent used to probe transcription elongation defects^27^ (Fig. 3h), underscoring its functional importance.

TL folding is also tightly coupled to BH bending (Figs 3a, 3d). From the pre-template-loading (Frame 1, State I) to the pre-catalysis state (Frame 20, State V), the BH bending edge (Ala828) shifts upward by ∼2.3 Å (Fig. 3d, top). A top-down view reveals opposite directional movements of BH segments (Thr809–Ala828 and Val829–Ile848), reflecting a pronounced bending motion (Fig. 3f, bottom). Early in the cycle (Frame 1-7), BH residue Lys830 inserts into the unfolded TL (Fig. 3f, left). During TL folding (Frame 8–10), BH rotation repositions Lys830 upward, avoiding steric clashes with TL residue Thr1080 (Fig. 3f, middle and right), further highlighting the coordinated rearrangements essential for catalysis.

### Substrate-induced RNAPII EC tightening

Conformational dynamics are not restricted to the active site but extend across the entire RNAPII EC through an interaction network that couples local rearrangements to global structural transitions (Fig. 4; Extended Data Fig. 8; Movie S5). To analyze these large-scale motions, we partitioned the EC into two structural modules: module 1, primarily composed of Rpb1 (Fig. 4a–b, light blue), and module 2, largely corresponding to Rpb2 (Fig. 4a–b, tan).

Module 1 and module 2 undergo opposing rotations around the RNA-DNA hybrid axis, with module 2 rotating counterclockwise (∼4°) and module 1 rotating clockwise (∼2°) from Frame 1 to Frame 20 (Fig. 4a, 4e). These rotations are coordinated with the conformational changes observed at the active site, including movements of the TL folding, BH bending and funnel closure (Fig. 4a). RNAPII adopts a pincer-like architecture in which module 1 and module 2 form two opposing arms, while the nucleic acid scaffold is positioned at the base of the cleft and clamped between them (Fig. 4a–b). Upon substrate binding, these two modules move toward each other along the downstream DNA tunnel, resulting in narrowing of the nucleic acid tunnel and formation of a more compact, closed conformation (Fig. 4a–c). We also found that the Rpb4/7 stalk moves in concert with module 1 in a coupled manner (Extended Data Fig. 8e), although in several intermediate states the stalk was too flexible to be modeled.

To quantitatively characterize this motion, we selected two representative domains from each module: the clamp domain from module 1 and the lobe domain from module 2, which are positioned on opposite sides of the downstream DNA tunnel. During clamp closure, the distance between these domains decreases from ∼24 Å to ∼17 Å (Fig. 4c), providing direct structural evidence for tunnel narrowing.

To elucidate the pathways underlying these global conformational changes, we mapped an interaction network that connects local rearrangements at the active site to distal regions of RNAPII (Extended Data Fig. 9a–b). Substrate binding induces structural changes within the active site that are propagated through this network, linking active-site elements with surrounding domains and reshaping the overall RNAPII conformation. Individual conformational events were resolved along the 3DVA trajectory and mapped onto the corresponding frame spectrum (Extended Data Fig. 9c), capturing the conformational continuum underlying the transition. Together, these results describe a substrate-induced, coordinated conformational transition of RNAPII that corresponds to progressive EC tightening.

Because Dataset 2 used a short RNA scaffold lacking upstream RNA, the observed tightening could potentially reflect the absence of upstream RNA rather than an intrinsic substrate-induced conformational change. To address this possibility, we reconstituted an RNAPII EC using an extended RNA scaffold containing a 19-mer RNA, designed to mimic a more mature elongation complex. The complex was assembled using the same strategy as in Dataset 2, with a 3′-dOH RNA primer in the presence of ATP substrate. From this dataset, referred to as Dataset 4, we resolved two structures corresponding to the TL-closed and TL-open states at resolutions of 2.5 Å and 2.6 Å, respectively (Extended Data Fig. 5; Table S5). The TL-closed state closely corresponds to Frame 20, State V, in Dataset 2, whereas the TL-open state resembles the substrate-bound state, Frame 7, State III.

Notably, clear density was observed for RNA extending into the exit tunnel, confirming proper assembly of a fully formed, mature elongation complex containing upstream RNA (Extended Data Fig. 10a). Structural comparison between the TL-open and TL-closed states revealed consistent conformational transitions, including TL folding, funnel closure, BH bending, clamp closure, and coordinated repositioning of the nucleic acid scaffold (Extended Data Fig. 10b–e). These observations demonstrate that substrate-induced EC tightening occurs in both short RNA scaffolds and mature elongation complexes containing upstream RNA.

### Nucleic acid scaffold repositioning couples to substrate alignment for catalysis

Accompanying clamp closure and global RNAPII tightening, the nucleic acid tunnel narrows, leading to a coordinated repositioning of the nucleic acid scaffold within the EC (Fig. 4d–e). These changes involve the non-template DNA strand (ntsDNA) of the transcription bubble (Fig. 4d), as well as the downstream and upstream DNA duplexes, and the RNA-DNA hybrid (Fig. 4e).

Within the transcription bubble, the ntsDNA region undergoes pronounced reorganization. As clamp closure proceeds, the lobe domain of module 2 shifts ∼5 Å upward and pushes against the ntsDNA, promoting progressive ordering of this region (Fig. 4d; Extended Data Fig. 8c). The downstream DNA duplex exhibits relatively modest movement, coordinated with the lower jaw domain of module 1, undergoing a ∼3° rotation and a ∼5 Å displacement (Fig. 4e; Extended Data Fig. 8e). In contrast, the upstream DNA shows pronounced rearrangement. This motion is coordinated with module 2 and involves direct interactions with the protrusion domain, wall domain, and Rpb12 subunit (Fig. 4d; Extended Data Fig. 8e). The upstream DNA undergoes an approximately ∼40° rotation and a ∼16 Å displacement, reflecting large-amplitude repositioning (Fig. 4e).

In contrast to other regions of the nucleic acid scaffold, the RNA-DNA hybrid undergoes relatively minor positional changes. However, these subtle adjustments are functionally important. From State I to State V (Frame 1 to Frame 20), the RNA 3′-terminal nucleotide progressively approaches the ATP substrate by ∼2 Å (Fig. 4e, right closed-up view; Extended Data Fig. 8d). In the pre-catalysis state (State V), a 3′-OH group was modeled onto the RNA primer, revealing a closest distance of 3.6 Å to the ATP α-phosphate, consistent with an optimal geometry for nucleophilic attack and coordination with the catalytic metal ions. These observations highlight that even small adjustments of the RNA-DNA hybrid are critical for achieving precise catalytic alignment and promoting efficient nucleotide incorporation.

### Post-catalysis structures and conformational states

Capturing RNAPII EC conformations immediately after phosphodiester bond formation has long been a challenge due to the rapid kinetics of nucleotide incorporation, leaving a critical gap in our understanding of the complete NAC. To overcome this, we developed a substrate-gradient strategy on cryo-EM affinity grids, enabling the capture of transient post-catalysis intermediates (see Methods for details). Using this approach, we established a native RNAPII EC assembled with an unmodified RNA primer and all four NTP substrates (ATP, UTP, CTP, GTP), allowing RNA extension. In this dataset (Dataset 3), we resolved five previously uncharacterized post-catalysis structures (States VI and VII-A–VII-D), in addition to the states observed in Dataset 2 (States II and III) (Fig. 5; Extended Data Figs. 3–4; Table S4).

The earliest post-catalysis intermediate, State VI, features a newly formed phosphodiester bond with the TL remaining in a closed conformation, representing the product state (Fig. 5a [VI], 5b; Movie S7). In this state, pyrophosphate (PPi) remains bound at the A-site, interacting with TL residue His1085 (Fig. 5b; Extended Data Fig. 11). Subsequent intermediates (States VII-A–VII-D) capture a stepwise TL opening and unfolding process (Fig. 5a [VII], 5c–f; Movie S8). In State VII-A, the TL tip (Phe1084–Ser1091) becomes disordered (Fig. 5c). In State VII-B, the TL helix (Lys1092–Val1107) continues to unfold, with the loop (Lys1092–Gly1097) re-engaging the cleft domain while the opposing helix (Met1063–His1085) begins to destabilize and His1085 disengages from PPi (Fig. 5d). In State VII-C, the TL segment Met1063–Leu1081 unfolds further, and Leu1081 releases from the RNA 3′-end base (Fig. 5e). State VII-D captures the fully open TL, coinciding with PPi relocation to the E-site, consistent with TL opening creating the clearance required for PPi release (Fig. 5f).

### Post-catalysis RNAPII EC relaxation

Coupled with PPi generation and TL opening in the active site, the RNAPII EC undergoes a coordinated, global transition toward a more relaxed state. This process represents the reverse of the tightening transition observed during the pre-catalysis phase (Dataset 2 and 4) and is therefore defined as “post-catalysis EC relaxation”. In Dataset 3, such a transition is captured through a series of structural intermediates spanning State VI to State VII (A–D) (Fig. 6a).

**Figure 6.**
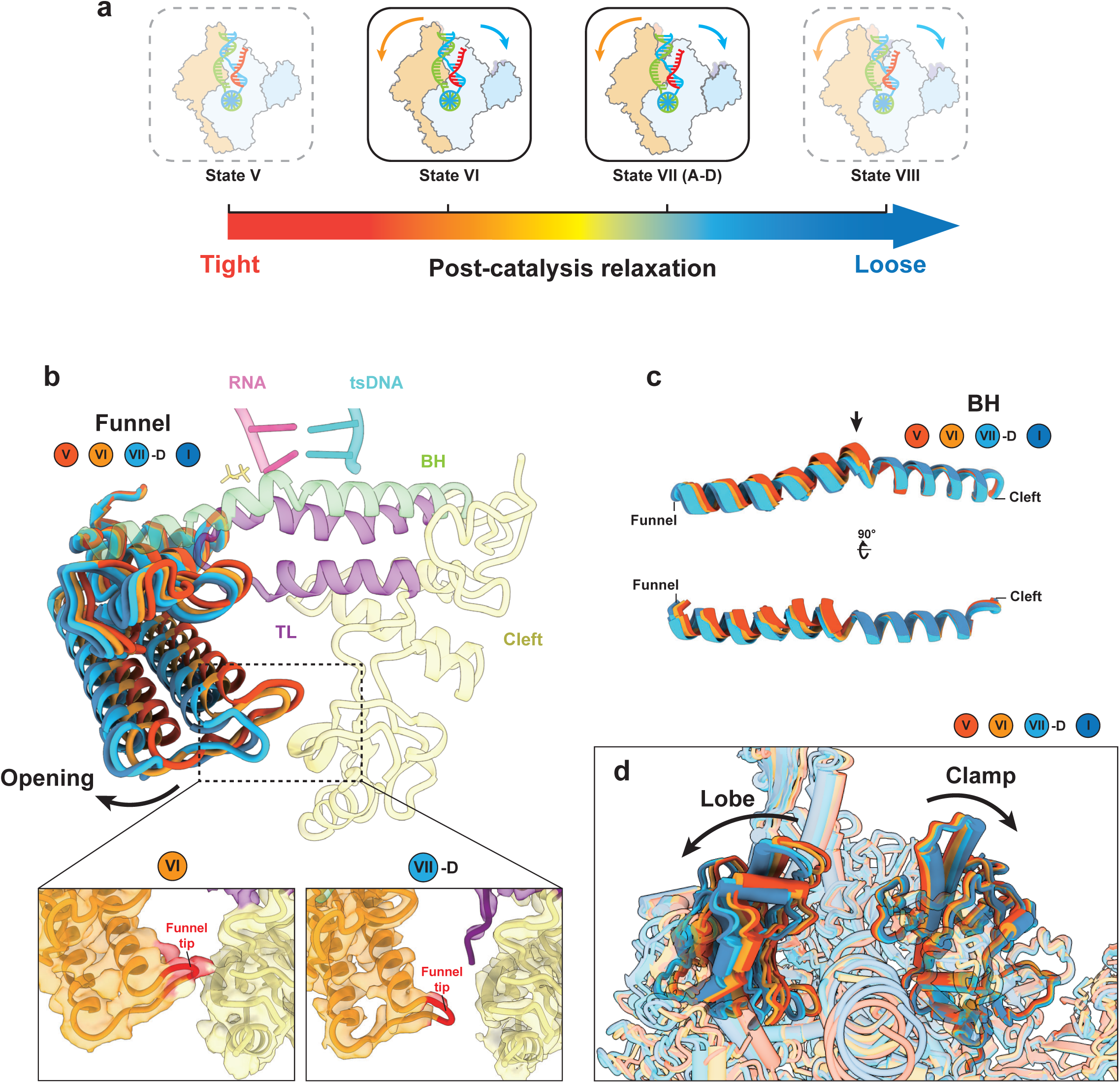
Post-catalysis RNAPII EC relaxation revealed by Dataset 3. (a) Schematic representation of the conformational spectrum of RNAPII EC in dataset 3. Four states (state V–VIII) are arranged along a linear spectrum, transitioning from a tight conformation (red, state V) to a loose conformation (blue, state VIII). Cartoon representations of RNAPII EC are shown for each state, with module 1 and module 2 colored in light blue and light orange, respectively. Arrows indicate the relative rotational movement between the two modules. States that were not resolved in Dataset 3 are shown as transparent and outlined by dashed lines, whereas states classified from Dataset 3 (state VI, and VII) are enclosed by solid outlines. (b–d) Structural comparison of local conformational changes across different states. Superposition of selected structures highlights conformational dynamics in the funnel helix (b), bridge helix (c), and lobe/clamp domain (d). Two states derived from the Dataset 3 (state VI, and VII) are compared together with reference states representing the extremes of the conformational spectrum from Dataset 2 (frame 1 at state I, loosest; frame 20 at state V, tightest). Color coding is consistent with conformational spectrum in panel a.

Structural comparisons across datasets illustrate this relaxation process. Specifically, State VI and State VII-D from Dataset 3 were analyzed alongside the reference states State V (tight) and State I (open) from Dataset 2 (Fig. 6b–d). State VI represents an immediate post-catalysis intermediate that remains in a tightly engaged conformation, closely resembling the pre-catalysis closed state (State V). In contrast, the State VII ensemble adopts progressively more open conformations, indicating a transition away from the catalytically competent state. This comparison reveals a coordinated global relaxation of the EC, accompanied by local structural rearrangements in multiple regions, including funnel opening, BH unbending, and clamp opening. These regional changes together define the transition of the EC from a compact, catalytically poised configuration toward a more relaxed and translocation-ready state (Fig. 6b–d). Notably, this conformational progression mirrors, in reverse, the tightening transition observed during substrate binding in Dataset 2 and 4, supporting a unified framework of catalysis-dependent conformational cycling.

## Discussion

### The complete NAC of RNAPII

We propose a complete NAC model for the RNAPII EC, comprising States I–VIII and including four previously uncharacterized intermediates: States I, V, VI, and VII (Fig. 7). These states delineate three major functional stages: translocation (shaded blue), substrate binding (shaded yellow), and catalysis (shaded orange), followed by post-catalysis reset.

**Figure 7.**
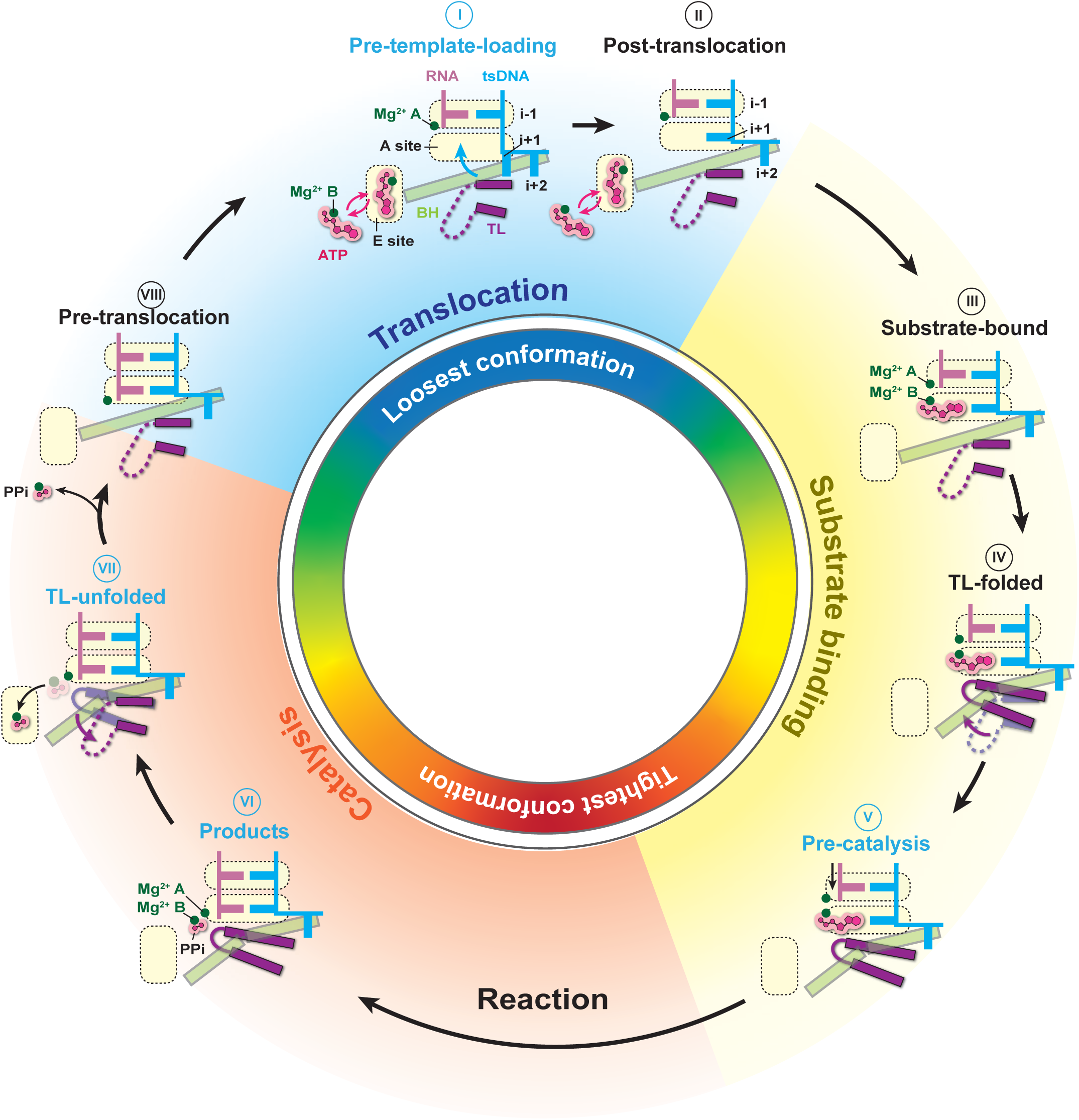
Comprehensive cycle of substrate incorporation by RNAPII EC. Schematic depicting the sequential transition through eight distinct conformational states (states I to VIII) within the active center of RNAPII during the nucleotide addition cycle. Newly resolved states in this study are highlighted with blue annotations. The complete nucleotide incorporation cycle is divided into three main phases, each denoted by a distinct background color: translocation (blue), substrate binding (yellow), and catalysis (orange). The inner cycle displays a color gradient representing clamp conformations ranging from loose (blue) to tight (red). Specific domains are color-coded for clarity: TL (purple), BH (light green), RNA primer (red), and template DNA (cyan).

The translocation stage (Fig. 7, shaded blue) comprises the pre-translocation state (State VIII), pre-template-loading state (State I), and post-translocation state (State II), which exist in dynamic equilibrium. During this phase, RNAPII adopts a relatively open and flexible conformation with an unfolded TL. The active site remains accessible, allowing incoming NTPs to diffuse nonspecifically into the E site in both States I and II.

The substrate-binding stage (Fig. 7, shaded yellow) begins when a cognate NTP base-pairs with the template at the *i*+1 position in the A site. In State III, the substrate is bound while the TL remains open. Substrate recognition initiates progressive TL folding (State IV), triggering conformational changes throughout the EC that culminates in the pre-catalysis state (State V). In this state, the TL is fully closed and the RNA 3′-OH is precisely aligned with the NTP α-phosphate, positioning the active site for catalysis.

Following phosphodiester bond formation, the complex enters the post-catalysis stage (Fig. 7, shaded orange). In the immediate product state (State VI), the TL remains closed, stabilizing the EC while metal-coordinated pyrophosphate (PPi) remains bound at the A site. The TL then unfolds (States VIIA–D), clearing space for PPi release. The cycle concludes with the pre-translocation state (State VIII), in which the TL is open, PPi has dissociated, and the active site is reset for the next round of nucleotide addition.

### Dynamic coordination of template loading during translocation

The pre-template-loading state (State I) identified here provides new insight into the temporal coordination of template loading and substrate binding within the NAC. This previously uncharacterized intermediate precedes the canonical post-translocation state (State II) and represents the earliest initiation step of a new cycle. Importantly, this state provides direct structural evidence that an NTP can associate with the E-site before the template base is loaded, offering a mechanistic explanation for kinetic observations that nucleotide binding to RNAPII can occur both before and after downstream DNA translocation^15,18^.

This pre-template-loading state differs from the half-translocation intermediate observed in a paused RNAPII EC by Cramer and colleagues^28^. In the paused structure, the RNA has shifted upstream, but the upstream tsDNA remains in a pre-translocation position. In contrast, our structure shows that both the RNA and upstream tsDNA have translocated, whereas the downstream template base has not yet entered the active site. These findings suggest that upstream and downstream translocation could occur in a partially unsynchronized manner, rather than as a concerted movement (at least in early elongation phase). We note that State I may be transient and difficult to capture, and that our experimental conditions, such as the use of a short RNA scaffold and/or a specific nucleic acid sequence, may have stabilized this state. Further studies will be needed to elucidate the dynamic properties of State I.

### A catalysis-associated conformational dynamic linking the active site to global RNAPII architecture

Our study reveals a previously unrecognized catalysis-associated conformational dynamic that extends from the active site to the entire RNAPII EC (Movie S1; Extended Data Fig. 9). While earlier studies established that NTP binding promotes TL folding and BH bending^12,29^, the accompanying global architectural changes in RNAPII have remained unclear. Here, we observe large-scale conformational dynamics consistently across multiple datasets, including short- and long-RNA scaffolds as well as native NTP-containing conditions, suggesting that this motion represents a general feature of RNAPII.

The global conformational dynamics of RNAPII can be divided into two coordinated phases: a substrate-induced EC tightening phase and a post-catalysis EC relaxation phase. During substrate engagement, conformational changes are initiated within the active site, where TL folding coordinates with funnel rotation and BH bending to form a dynamic core. This core is integrated with the rest of RNAPII through an interaction network, thereby spreading local active-site rearrangements into global structural changes across the complex. At the global level, RNAPII behaves as a pincer-like assembly, with module 1 and module 2 forming two opposing arms that enclose the nucleic acid scaffold at the base of the cleft. Upon substrate loading, these modules undergo coordinated rotations toward the downstream DNA channel, resulting in clamp closure and narrowing of the nucleic acid tunnel. This global tightening is accompanied by repositioning the nucleic acid scaffold, improving alignment of the RNA primer 3′ terminus with the incoming substrate and establishing a catalytically competent configuration.

This RNAPII tightening is distinct from the swiveling motion described in bacterial RNA polymerase (RNAP) (Extended Data Fig. 12). Bacterial swiveling is primarily associated with transcriptional pausing and translocation-related intermediates^30,31^, where the swivel module rotates approximately parallel to the nucleic acid scaffold (Extended Data Fig. 12a,b). By contrast, the tightening observed here is reproducibly detected upon substrate binding across Datasets 2–4, but not in the substrate-free Dataset 1, supporting its interpretation as substrate-induced EC tightening. Structurally, this motion involves coordinated movement of RNAPII modules 1 and 2 toward the nucleic acid scaffold, narrowing the cleft in a pincer-like manner (Extended Data Fig. 12c).

Following phosphodiester bond formation, the TL unfolds, accompanied by reopening of the funnel domain, straightening of the BH, and reversal of clamp closure. These changes widen the nucleic acid tunnel and restore RNAPII to a more open, relaxed conformation. We refer to this process as post-catalysis relaxation.

Together, these two phases define a cyclic conformational mechanism in which RNAPII transits between a tightly engaged, reaction-ready state (red/orange) and a relaxed, translocation-competent state (blue/green) (Fig. 7 inner cycle; Movie S1). This switching behavior, resembling a molecular spring, links active-site chemistry to global structural transitions and may represent a general mechanism for allosteric regulation in multi-subunit polymerases.

## Methods and Materials

### RNAPII purification and elongation complex assembly

*Saccharomyces cerevisiae* 12-subunit RNA polymerase II (RNAPII) was purified and assembled *in vitro* as described previously^12,32^. The 12-subunit RNAPII was further assembled with biotinylated nucleic acid scaffold to generate the RNAPII elongation complex (EC) as previously reported^16^. All RNA and DNA oligonucleotides used for nucleic acid scaffolds were purchased from Integrated DNA Technologies (IDT). The nucleic acid sequences are follows: Biotinylated template strand DNA (tsDNA): 5′-/BiotinTEG/ TTT TTT GAT ATT TTT GGA TCC CGC TCT GCT CCT TCT CCC ATC CTC TCG ATG GCT ATG AGA TCA ACT AGG AAT TC-3′; Biotinylated non-template strand DNA (ntsDNA): 5′-/BiotinTEG/ TTT TTA TGT ATT AAT GAA TTC CTA GTT GAT CTC ATA GCC CAT TCC TAC TTG GGA GAA GGA GCA GAG CGG GAT CC-3′; RNA (9-mer): 5′-AUC GAG AGG-3′; RNA (19-mer): 5′-UUU UUU UUU CAU CGA GAG G-3′. Notably, a 3’-deoxy RNA, instead of a regular RNA, was used in the sample of Dataset 1, 2, and 4.

### Cryo-EM sample preparation using mspSA affinity grids

We employed mspSA affinity grids for cryo-EM data collection, which effectively overcome the orientation bias and air-water interface issue of RNAPII and improved resolution^16^. The optimized protocol for preparing mspSA affinity grids is outlined as follows: Mix DOPC (18:1 (Δ9-Cis) PC, Avanti) and biotin-labeled lipid (16:0 Biotinyl Cap PE, Avanti) in a 9:1 ratio to reach a final concentration of 1 mg/mL. Prepare a streptavidin solution (streptavidin from Streptomyces avidinii, Sigma) at a concentration of 0.01 mg/mL in a buffer consisting of 20 mM HEPES (pH 7.5) and 150 mM NaCl. Add 30 µL of the streptavidin solution to a Teflon well (on ice), and then carefully deposit 1 µL of the lipid mixture onto the surface of the streptavidin solution. Enclose the Teflon block in a humidity chamber containing a small amount of Milli-Q water at the bottom of chamber. Seal the lid of the chamber with Vaseline and incubate overnight at room temperature. On the following day, carefully place the carbon side of holy carbon grids (Quantifoil copper 2/1, 300 mesh) onto the surface of the Teflon well (without prior glow discharge) for 2 minutes to allow for lipid monolayer transfer. Wash the grid with three drops (120 µL) of sample buffer to remove unbound streptavidin. Add 4 µL of the RNAPII EC samples to affinity grids and incubate for approximately 5 minutes. Subsequently, plunge-freeze the grids using the Thermo Fisher Vitrobot IV, applying the following parameters: 4 seconds blotting time, −10 blotting force, at 4°C and 100% humidity with adding 4 µL of sample buffer. The blotting force should be optimized to achieve the optimal vitrification results.

Four samples with different ECs and substrates were prepared for cryo-EM data collection, including the apo EC containing 9-mer 3’-deoxy RNA (Dataset 1), EC containing 3’-deoxy 9-mer RNA with substrate ATP (Dataset 2), EC containing native 9-mer RNA with substrate NTP (Dataset 3), and EC containing 3’-deoxy 19-mer RNA with substrate ATP (Dataset 4) (Extended Data Figs. 1, 2, 3, and 5). For Dataset 1, no substrate was added and RNAPII EC was directly applied to the affinity grids. For Dataset 2 and 4, 2 mM substrate ATP and RNAPII EC were pre-mixed for 5 mins at room temperature prior to application onto the affinity grids.

For Dataset 3, to capture transient elongation states during active nucleotide addition, we employed a substrate-gradient affinity grid (SGAG) strategy. Specifically, biotin-tagged RNAPII ECs were first applied to streptavidin-coated affinity grids and incubated for 5 minutes to allow stable immobilization. Unbound ECs were subsequently removed by washing twice gently with 50 μL of fresh buffer, and excess buffer was gently absorbed from the edge of the grid using filter paper. The grid was then transferred to a Thermo Fisher Vitrobot IV equilibrated at 4 °C and 100% humidity. Before blotting, 3 μL of fresh buffer was added to the grid surface, followed by application of a small droplet (0.2 μL) of a highly concentrated NTP mixture (10 mM total, containing all four NTPs: ATP, UTP, CTP, and GTP) from the upper edge of grid. The grid was immediately blotted for 1 second and plunge frozen. During this brief interval, NTPs diffuse and generate a transient concentration gradient across the grid (Extended Data Fig. 3). Sampling across this NTP concentration gradient enables capture of a range of potentially transient intermediate transcriptional states of ECs within the time frame of vitrification (Extended Data Fig. 3).

### Data collection, processing and 3D reconstruction

Initial screening to obtain grids with well-formed monolayers and particles was conducted using a 200 kV Thermo Scientific™ Glacios™ transmission electron microscope. Cryo-EM data were collected using a FEI Titan Krios operating at 300 kV, equipped with a Gatan K3 with GIF Quantum camera (at eBIC, Diamond). Data acquisition was performed automatically using EPU software, with defocus values ranging from −0.5 to −2.5 µm. The imaging pixel size for data collection was 0.829 Å (Bio-Quantum K3) for Dataset 1 and Dataset 2, 0.825 Å for Dataset 3, and 0.932 Å for Dataset 4. The total dose applied was 50 electrons per Å² for all samples distributed across 50 frames and Dataset 4 was recorded in EER format. A total of 15,165, 12,739, 18,913, 2000 images were collected for Dataset 1–4, respectively.

Data were processed via CryoSPARC (v4.5.3)^33^. Datasets 1–4 were analysed using a similar pipeline, as described below. Raw micrographs were imported, followed by motion correction and calculation of the contrast transfer function (CTF). The resulting micrographs were manually curated to exclude images with poor quality, (such as CTF fit resolution worse than 5 Å; Relative ice thickness: 1< values < 1.1; Total full-frame motion distance (pixels): < 60 and astigmatism: < 1000). All micrographs were then subjected to automatic particle picking (Blob picking), with particle diameters ranging from 150 Å to 190 Å. Following several rounds of 2D classification, particles displaying high-quality 2D class averages were selected for *ab initio* reconstruction, followed by heterogeneous refinement. Particles from well-resolved 3D classes obtained after heterogeneous refinement were used for Topaz training for picking^34,35^. After one or more additional rounds of Topaz training, the resulting high-quality particles were subjected to *ab initio* reconstruction and further heterogeneous refinement. Particles from well-defined 3D classes, together with high-quality particles obtained through blob picking, were combined, and duplicate entries were removed. The refined particle set was then used for another round of *ab initio* reconstruction and heterogeneous refinement, followed by homogeneous refinement with defocus refinement and global CTF refinement, enabling the best 3D class obtained from heterogeneous refinement. To separate distinct conformational states across all three datasets, 3D classification with a solvent mask was performed without alignment. For Dataset 2, 3D Variability Analysis (3DVA)^25^ were conducted in intermediate mode, in which all particles were subdivided into 20 groups and further refined using homogeneous refinement. In all cases, the final resolution was assessed using the gold-standard Fourier shell correlation (FSC) criterion. Local resolution estimation was performed in UCSF Chimera^36^ and ChimeraX^37^ using the output maps generated by CryoSPARC.

In parallel, Dataset 2 was subjected to conventional 3D classification employing a solvent mask and omitting alignment procedures. This analysis yielded structural ensembles that delineate a coherent trajectory of temporal conformational transitions—progressing from an open TL to a fully closed state with substrate bound—remarkably concordant with those resolved via 3D Variability Analysis (3DVA) under both simple and intermediate modes (Extended Data Fig. 6). The convergence of these independent analytical modalities not only reinforces the robustness of our findings but also affirms the reproducibility and fidelity of the captured intermediate conformational states.

### Atomic model building, refinement, and validation

We generated a total of 31 atomic models of RNAPII EC from four cryo-EM datasets. Specifically, the Dataset 1 yielded two distinct structures representing the pre-template-loading state (State I) and the post-translocation state (State II). From Dataset 2, we resolved 20 structural intermediates of RNAPII EC, covering dynamic transitions corresponding to State I through V, aligned with the principal component analysis (PC0) results of 3DVA. From the Dataset 3, seven structures were determined, comprising one product state (State VI), four sequential TL opening intermediates (States VII-A to VII-D), one post-translocation state (State II) and one substrate-bound state (State III). For the Dataset 4, two structures were determined at State III (TL-open) and V (TL-closed).

For all these models, we used the same pipeline for model building, refinement, and validation. The previously published RNAPII EC structure (PDB: 2E2H^12^ and 8U9R^13^) was used as the initial model. The resulting models were rigid-body fitted into the cryo-EM density maps using ChimeraX UCSF (v1.8)^36^. Model refinement was performed in PHENIX (v1.21)^38^ using the phenix.real_space_refine program, followed by manual adjustments in COOT (v0.9.8)^39^. Model validation was conducted with MolProbity^40^. All structural figures were rendered using ChimeraX UCSF (v1.8)^36^.

### MPA sensitivity assay

The funnel tip deletion (Rpb1 H706–M708 deletion) strain was generated in the BJ5465 background (*MAT*a *ura3-52 trp-1 leu2*_*1 his3*_*200 pep4*::*HIS3 prb1*_*1.6R can1*)^41^ using CRISPR/Cas9. The deletion was confirmed by DNA sequencing.

For the MPA sensitivity assay, wild-type and Rpb1 funnel tip-deleted strains were grown in yeast extract peptone dextrose (YPD) medium at 30°C to stationary phase. Cultures were then serially diluted 10-fold, and 5 µl of each dilution was spotted onto YPD plates supplemented with the indicated concentrations of MPA. Plates were incubated at 30°C for 4 days before imaging.

### *In vitro* transcription assay

RNAPII EC samples were assembled using the nucleic acid scaffold (containing regular 9-mer RNA) identical to the one used for cryo-EM sample preparation, with slight modifications in complex assembly process. Initially, the radioactively labeled RNA primer (regular 9-mer) was annealed with the template DNA (tsDNA). Subsequently, the 12-subunit RNAPII was incubated with the RNA-tsDNA hybrid, followed by the addition of non-template strand DNA (ntsDNA) to complete the assembly of the RNAPII EC. The final ratio of components in the RNAPII EC was as follows: RNAPII: RNA: tsDNA: ntsDNA = 6:1:2:3. RNAPII EC samples were prepared at a final concentration of 40 nM in elongation complex (EC) buffer containing 20 mM Tris-HCl (pH 7.5), 5 mM MgCl_2_, 40 mM KCl, and 5 mM DTT. *In vitro* transcription was initiated by the addition of an NTP mixture. Three substrate NTP concentrations (5 mM, 1 mM, and 200 μM) were tested here. For each concentration, transcription reactions were terminated at multiple time points: 5 s, 10 s, 30 s, and 10 min. Reactions were quenched by the addition of an equal volume of quench-loading buffer (90% formamide, 50 mM EDTA, 0.05% xylene cyanol, and 0.05% bromophenol blue), followed by heating at 95 °C for 10 min. Reaction products were analyzed by denaturing polyacrylamide gel electrophoresis (PAGE) containing 6 M urea. The gel was visualized by phosphorimaging, and bands were quantified using Image Laboratory software (Bio-Rad).

## Data availability

All data needed to evaluate the conclusions in the paper are present in the paper and/or the supplementary information. Source Data are provided with this paper. Cryo-EM density maps and corresponding atomic coordinates have been deposited in the Electron Microscopy Data Bank (EMDB) and Protein Data Bank (PDB), respectively. Accession codes are as follows: Maps and/or atomic models of RNAPII EC-Apo (Dataset 1) have been deposited under accession code for State-I (EMD-54704/PDB 9SAY), State**-**I-A (EMD-54705), State-I-B (EMD-54706) and State-II (EMD-54707/PDB 9SAZ). Accession codes of maps and atomic models of RNAPII EC in complex with ATP (Dataset 2) generated from 3D variability and 3D display (Intermediate model, Window=0) are EMD-54730/ PDB 9SBL, EMD-54731/ PDB 9SBM, EMD-54732/ PDB 9SBN, EMD-54733/ PDB 9SBO, EMD-54734/ PDB 9SBP, EMD-54735/ PDB 9SBQ, EMD-54736/ PDB 9SBR, EMD-54737/ PDB 9SBS, EMD-54738/ PDB 9SBT, EMD-54739/ PDB 9SBU, EMD-54740/ PDB 9SBV, EMD-54741/ PDB 9SBW, EMD-54742/ PDB 9SBX, EMD-54743/ PDB 9SBY, EMD-54744/ PDB 9SBZ, EMD-54745/ PDB 9SC0, EMD-54746/ PDB 9SC1, EMD-54747/ PDB 9SC2, EMD-54748/ PDB 9SC3 and EMD-54749/ PDB 9SC4 (ordered from Frame1 to Frame 20). Additional maps obtained from 3D classification (with solvent mask) have been deposited separately: 3D Class A (Frame 1) (EMD-54677), 3D Class B (Frame 4) (EMD-54682), 3D Class C (Frame 7) (EMD-54681), 3D Class D (Frame 8) (EMD-54680), 3D Class E (Frame 9) (EMD-54679), 3D Class F (Frame 10) (EMD-54683), 3D Class G (Frame 13) (EMD-54684), 3D Class H (Frame 15) (EMD-54685), 3D Class I (Frame 17) (EMD-54686), and 3D Class J (Frame 20) (EMD-54687). Maps and atomic models of RNAPII EC in complex with NTP (Dataset 3) have been deposited under accession code for State-II (EMD-54713/9SB5), State-III (EMD-57322/29RF), State-VI (EMD-54708/ PDB 9SB0), State-VII-A (EMD-54709/ PDB 9SB1), State-VII-B (EMD-54710/ PDB 9SB2), State-VII-C (EMD-54711/ PDB 9SB3), and State-VII-D (EMD-54712/ PDB 9SB4). Accession codes of maps and atomic models of long-RNA EC in complex with ATP (Dataset 4) are State III (TL-open) (EMD-57673/PDB 30EP) and State-V (TL-closed) (EMD-57676/PDB 30ES).

## Acknowledgements

This work was supported by grants from the National Institutes of Health (R01 GM102362 and GM148476 to D.W.; U54 AI170791-7522 and R21 CA280467 and R21 AI184080 to P.Z.), the UK Wellcome Trust Investigator Award (206422/Z/17/Z to P.Z.), the UK Wellcome Discovery Award (311427/Z/24/Z to P.Z.), ERC AdG grant (101021133 to P.Z.), and the National Science Foundation (MCB-2102072 to S.L.). We thank Diamond Light source for access and support of the cryoEM facilities at the UK National Electron Bio-Imaging Centre (eBIC) (proposal NT29812, NR21005). Further electron microscopy provision was provided through the OPIC electron microscopy facility, a UK Instruct-ERIC Centre, which was founded by a Wellcome JIF award (060208/Z/00/Z) and is supported by a Wellcome equipment grant (093305/Z/10/Z). Computation was performed at Diamond Light Source, the Oxford Biomedical Research Computing (BMRC) facility, a joint development between the Wellcome Centre for Human Genetics (Wellcome Trust Core Award Grant Number 203141/Z/16/Z) and the Big Data Institute (BDI) supported by Health Data Research UK and the NIHR Oxford Biomedical Research Centre.

## Author contributions

D.W. and P.Z. conceived and supervised the project. Q.L. performed protein purification. G.Y. prepared the cryo-EM samples on mspSA affinity grids and performed cryo-EM data collection, data processing and cryo-EM SPA map reconstruction under the supervision of P.Z. D.C. provided guidance and support for data processing. Q.L. and G.Y. built and validated the structural models, and analysis under the supervision of P.Z. and D.W. H.H. and S. L. created the funnel tip mutant and performed functional analysis of the mutant. Q.L., G.Y., P.Z. and D.W. performed data analysis. Q.L., G.Y., and D.W. made the figures. Q.L., G.Y., P.Z. and D.W. wrote the manuscript, with input from all authors.

