## Supplementary Tables and Figures for "Structural Dynamics of RNA Polymerase II During Nucleotide Addition Cycle"

**Table S1 | Cryo-EM data collection and structure determination of RNAPII EC without substrate (Dataset 1)**

| Map | State-I | State-I-A | State-I-B | State-II |
|---|---|---|---|---|
| **PDB** | 9SAY | NA | NA | 9SAZ |
| **EMDB** | EMD-54704 | EMD-54705 | EMD-54706 | EMD-54707 |
| **Data collection** | | | | |
| Microscope | Titan Krios Gatan | | | |
| Detector | K3 with GIF Quantum | | | |
| Cs (mm) | 2.7 | 2.7 | 2.7 | 2.7 |
| Magnification | 105k | 105k | 105k | 105k |
| Pixel size (Å) | 0.829 | 0.829 | 0.829 | 0.829 |
| Electron dose (e-/Å$^2$) | 50 | 50 | 50 | 50 |
| Defocus range (µm) | -0.5 to -2.5 | -0.5 to -2.5 | -0.5 to -2.5 | -0.5 to -2.5 |
| Micrograph Number | 15165 | 15165 | 15165 | 15165 |
| **Reconstruction** | | | | |
| Software | cryoSPARC-4.5 | | | |
| Particles refinement | cryoSPARC-4.5 | cryoSPARC-4.5 | cryoSPARC-4.5 | cryoSPARC-4.5 |
| Symmetry | 64243 | 57585 | 56940 | 72426 |
| Resolution (Å) | C1 | C1 | C1 | C1 |
| Sharpening B-factor (Å2) | 3.4 | 3.6 | 3.6 | 3.3 |
| **Refinement** | | | | |
| **Model composition** | | | | |
| Number of atoms | 32307 | NA | NA | 32314 |
| Protein residues | 3895 | NA | NA | 3896 |
| Nucleotides | 65 | NA | NA | 65 |
| Ligand | 8 ZN; 1 MG; | NA | NA | 1 MG; 8 ZN |
| **Bonds RMSD** | | | | |
| Bonds lengths (Å) | 0.003 | NA | NA | 0.0023 |
| Bonds angles (°) | 0.50 | NA | NA | 0.49 |
| **Validation** | | | | |
| MolProbity score | 1.71 | NA | NA | 1.58 |
| Clash score | 8.66 | NA | NA | 7.01 |
| Rotamer outliers | 0.00% | NA | NA | 0.15% |
| C-beta outliers | 0.00% | NA | NA | 0.00% |
| **Ramachandran plot** | | | | |
| Favored (%) | 96.35% | NA | NA | 96.82% |
| Allowed (%) | 3.55% | NA | NA | 3.13% |
| Outlier (%) | 0.10% | NA | NA | 0.05% |

### Table S2 | Cryo-EM data collection and structure determination of RNAPII EC with ATP substrate, and 3D Varibility Analyses (Dataset 2)

| **Map** | Frame 1 | Frame 2 | Frame 3 | Frame 4 | Frame 5 | Frame 6 | Frame 7 | Frame 8 | Frame 9 | Frame 10 | Frame 11 | Frame 12 | Frame 13 | Frame 14 | Frame 15 | Frame 16 | Frame 17 | Frame 18 | Frame 19 | Frame 20 |
|---|---|---|---|---|---|---|---|---|---|---|---|---|---|---|---|---|---|---|---|---|
| **PDB** | 9SBL | 9SBM | 9SBN | 9SBO | 9SBP | 9SBQ | 9SBR | 9SBS | 9SBT | 9SBU | 9SBV | 9SBW | 9SBX | 9SBY | 9SBZ | 9SC0 | 9SC1 | 9SC2 | 9SC3 | 9SC4 |
| **EMDB** | EMD-54730 | EMD-54731 | EMD-54732 | EMD-54733 | EMD-54734 | EMD-54735 | EMD-54736 | EMD-54737 | EMD-54738 | EMD-54739 | EMD-54740 | EMD-54741 | EMD-54742 | EMD-54743 | EMD-54744 | EMD-54745 | EMD-54746 | EMD-54747 | EMD-54748 | EMD-54749 |
| **Data collection** | | | | | | | | | | | | | | | | | | | | |
| Microscope | | | | | | | | | | Titan Krios Gatan | | | | | | | | | | |
| Detector | | | | | | | | | | K3 with GIF Quantum | | | | | | | | | | |
| Cs (mm) | | | | | | | | | | 2.7 | | | | | | | | | | |
| Magnification | | | | | | | | | | 105k | | | | | | | | | | |
| Pixel size (Å) | | | | | | | | | | 0.829 | | | | | | | | | | |
| Electron dose (e-/Å$^2$) | | | | | | | | | | 50 | | | | | | | | | | |
| Defocus range (μm) | | | | | | | | | | -0.5 to -2.5 | | | | | | | | | | |
| Micrograph Number | | | | | | | | | | 12,739 | | | | | | | | | | |
| **Reconstruction** | | | | | | | | | | | | | | | | | | | | |
| Software | | | | | | | | | | cryoSPARC-4.5 | | | | | | | | | | |
| Particles refinement | 16,262 | 22,773 | 29,765 | 37,985 | 46,826 | 56,496 | 66,397 | 77,148 | 86,647 | 94,160 | 98,766 | 100,314 | 97,126 | 89,544 | 79,845 | 67,276 | 53,389 | 40,178 | 29,059 | 19,500 |
| Symmetry | C1 | C1 | C1 | C1 | C1 | C1 | C1 | C1 | C1 | C1 | C1 | C1 | C1 | C1 | C1 | C1 | C1 | C1 | C1 | C1 |
| Resolution (Å) | 4.3 | 4.2 | 4.0 | 4.0 | 4.0 | 3.9 | 4.0 | 3.9 | 3.9 | 3.8 | 3.7 | 3.6 | 3.5 | 3.4 | 3.4 | 3.3 | 3.5 | 3.5 | 3.7 | 3.8 |
| Sharpening B-factor (Å$^2$) | 47.2 | 56.6 | 57.3 | 64.5 | 64.2 | 70.00 | 69.9 | 68.5 | 79.6 | 93.0 | 71.9 | 78.70 | 78.9 | 75.0 | 66.7 | 64.5 | 62.9 | 64.10 | 57.7 | 43.6 |
| **Refinement** | | | | | | | | | | | | | | | | | | | | |
| **Model composition** | | | | | | | | | | | | | | | | | | | | |
| Number of atoms | 32888 | 32901 | 32901 | 32901 | 32901 | 32895 | 32914 | 32965 | 32987 | 33083 | 33083 | 33083 | 33083 | 33083 | 33083 | 33083 | 33083 | 33083 | 33083 | 33083 |
| Protein residues | 3895 | 3897 | 3897 | 3897 | 3897 | 3896 | 3896 | 3902 | 3905 | 3908 | 3908 | 3908 | 3908 | 3908 | 3908 | 3908 | 3908 | 3908 | 3908 | 3908 |
| Nucleotides | 93 | 93 | 93 | 93 | 93 | 93 | 93 | 93 | 93 | 97 | 97 | 97 | 97 | 97 | 97 | 97 | 97 | 97 | 97 | 97 |
| Ligand | 8 ZN; 1 MG; 1 ATP | 8 ZN; 1 MG; 1 ATP | 8 ZN; 1 MG; 1 ATP | 8 ZN; 1 MG; 1 ATP | 8 ZN; 1 MG; 1 ATP | 8 ZN; 1 MG; 1 ATP | 8 ZN; 2 MG; 1 ATP | 8 ZN; 2 MG; 1 ATP | 8 ZN; 2 MG; 1 ATP | 8 ZN; 2 MG; 1 ATP | 8 ZN; 2 MG; 1 ATP | 8 ZN; 2 MG; 1 ATP | 8 ZN; 2 MG; 1 ATP | 8 ZN; 2 MG; 1 ATP | 8 ZN; 2 MG; 1 ATP | 8 ZN; 2 MG; 1 ATP | 8 ZN; 2 MG; 1 ATP | 8 ZN; 2 MG; 1 ATP | 8 ZN; 2 MG; 1 ATP | 8 ZN; 2 MG; 1 ATP |
| **Bonds RMSD** | | | | | | | | | | | | | | | | | | | | |
| Bonds lengths (Å) | 0.003 | 0.004 | 0.003 | 0.003 | 0.004 | 0.004 | 0.023 | 0.023 | 0.023 | 0.023 | 0.023 | 0.023 | 0.023 | 0.024 | 0.023 | 0.023 | 0.023 | 0.023 | 0.023 | 0.023 |
| Bonds angles (°) | 0.60 | 0.61 | 0.56 | 0.57 | 0.60 | 0.59 | 0.95 | 0.93 | 0.94 | 0.95 | 0.92 | 0.92 | 0.94 | 0.97 | 0.92 | 0.89 | 0.93 | 0.92 | 0.93 | 0.93 |
| **Validation** | | | | | | | | | | | | | | | | | | | | |
| MolProbity score | 1.90 | 1.95 | 1.80 | 1.71 | 1.86 | 1.89 | 1.84 | 1.77 | 1.81 | 1.89 | 1.68 | 1.63 | 1.75 | 1.70 | 1.63 | 1.63 | 1.65 | 1.62 | 1.71 | 1.74 |
| Clash score | 12.98 | 12.79 | 10.31 | 9.37 | 10.08 | 10.36 | 10.04 | 8.97 | 9.84 | 10.47 | 8.38 | 7.30 | 8.59 | 7.67 | 6.73 | 6.83 | 7.00 | 7.64 | 7.95 | 8.93 |
| Rotamer outliers | 0.03% | 0.03% | 0.00% | 0.00% | 0.03% | 0.00% | 0.00% | 0.00% | 0.00% | 0.20% | 0.00% | 0.00% | 0.00% | 0.00% | 0.00% | 0.00% | 0.00% | 0.00% | 0.00% | 0.00% |
| C-beta outliers | 0.00% | 0.00% | 0.00% | 0.00% | 0.00% | 0.00% | 0.00% | 0.00% | 0.00% | 0.00% | 0.00% | 0.00% | 0.00% | 0.00% | 0.00% | 0.00% | 0.00% | 0.00% | 0.00% | 0.00% |
| **Ramachandran plot** | | | | | | | | | | | | | | | | | | | | |
| Favored (%) | 95.98% | 95.20% | 96.07% | 96.61% | 95.23% | 94.92% | 95.41% | 95.78% | 95.81% | 94.96% | 96.52% | 96.47% | 95.85% | 95.90% | 96.24% | 96.24% | 96.08% | 96.73% | 95.93% | 96.13% |
| Allowed (%) | 3.97% | 4.64% | 3.88% | 3.34% | 4.72% | 5.00% | 4.49% | 4.09% | 4.14% | 4.88% | 3.38% | 3.43% | 3.99% | 4.00% | 3.66% | 3.66% | 3.79% | 3.14% | 3.55% | 3.55% |
| Outlier (%) | 0.05% | 0.16% | 0.05% | 0.05% | 0.05% | 0.08% | 0.10% | 0.13% | 0.05% | 0.16% | 0.10% | 0.10% | 0.16% | 0.10% | 0.10% | 0.10% | 0.13% | 0.13% | 0.10% | 0.13% |

#### Table S3 | Cryo-EM data collection and structure determination of RNAPII EC with ATP substrate, and 3D Classification Analyses (Dataset 2)

| **Map** | 3D-Class-A | 3D-Class-B | 3D-Class-C | 3D-Class-D | 3D-Class-E | 3D-Class-F | 3D-Class-G | 3D-Class-H | 3D-Class-I | 3D-Class-J |
|---|---|---|---|---|---|---|---|---|---|---|
| **EMDB** | EMD-54677 | EMD-54682 | EMD-54681 | EMD-54680 | EMD-54679 | EMD-54683 | EMD-54684 | EMD-54685 | EMD-54686 | EMD-54687 |
| **Data collection** | | | | | | | | | | |
| Microscope | Titan Krios Gatan | | | | | | | | | |
| Detector | K3 with GIF Quantum | | | | | | | | | |
| Cs (mm) | 2.7 | 2.7 | 2.7 | 2.7 | 2.7 | 2.7 | 2.7 | 2.7 | 2.7 | 2.7 |
| Magnification | 105k | 105k | 105k | 105k | 105k | 105k | 105k | 105k | 105k | 105k |
| Pixel size (Å) | 0.829 | 0.829 | 0.829 | 0.829 | 0.829 | 0.829 | 0.829 | 0.829 | 0.829 | 0.829 |
| Electron dose (e-/Å$^2$) | 50 | 50 | 50 | 50 | 50 | 50 | 50 | 50 | 50 | 50 |
| Defocus range (µm) | -0.5 to -2.5 | -0.5 to -2.5 | -0.5 to -2.5 | -0.5 to -2.5 | -0.5 to -2.5 | -0.5 to -2.5 | -0.5 to -2.5 | -0.5 to -2.5 | -0.5 to -2.5 | -0.5 to -2.5 |
| Micrograph Number | 12739 | 12739 | 12739 | 12739 | 12739 | 12739 | 12739 | 12739 | 12739 | 12739 |
| **Reconstruction** | | | | | | | | | | |
| Software | cryoSPARC-4.5 | cryoSPARC-4.5 | cryoSPARC-4.5 | cryoSPARC-4.5 | cryoSPARC-4.5 | cryoSPARC-4.5 | cryoSPARC-4.5 | cryoSPARC-4.5 | cryoSPARC-4.5 | cryoSPARC-4.5 |
| Particles refinement | 65010 | 73900 | 66560 | 93475 | 86363 | 44115 | 36095 | 69690 | 65841 | 88302 |
| Symmetry | C1 | C1 | C1 | C1 | C1 | C1 | C1 | C1 | C1 | C1 |
| Resolution (Å) | 3.6 | 3.6 | 3.5 | 3.5 | 3.7 | 4.0 | 4.1 | 3.3 | 3.3 | 3.2 |
| Sharpening B-factor (Å$^2$) | 72.9 | 67.7 | 73.3 | 80.3 | 77.6 | 62.9 | 57.4 | 68.4 | 64.5 | 68.2 |

**Table S4 | Cryo-EM data collection and structure determination of RNAPII EC with NTP substrate (Dataset 3)**

| **Map** | State-II | State-III | State-VI | State-VII-A | State-VII-B | State-VII-C | State-VII-D |
|---|---|---|---|---|---|---|---|
| **PDB** | 9SB5 | 29RF | 9SB0 | 9SB1 | 9SB2 | 9SB3 | 9SB4 |
| **EMDB** | EMD-54713 | EMD-57322 | EMD-54708 | EMD-54709 | EMD-54710 | EMD-54711 | EMD-54712 |
| **Data collection** | | | | | | | |
| Microscope | | | | Titan Krios Gatan | | | |
| Detector | | | | K3 with GIF Quantum | | | |
| Cs (mm) | 2.7 | 2.7 | 2.7 | 2.7 | 2.7 | 2.7 | 2.7 |
| Magnification | 105k | 105k | 105k | 105k | 105k | 105k | 105k |
| Pixel size (Å) | 0.825 | 0.825 | 0.825 | 0.825 | 0.825 | 0.825 | 0.825 |
| Electron dose (e-/Å$^2$) | 50 | 50 | 50 | 50 | 50 | 50 | 50 |
| Defocus range (μm) | -0.5 to -2.5 | -0.5 to -2.5 | -0.5 to -2.5 | -0.5 to -2.5 | -0.5 to -2.5 | -0.5 to -2.5 | -0.5 to -2.5 |
| Micrograph Number | 18913 | 18913 | 18913 | 18913 | 18913 | 18913 | 18913 |
| **Reconstruction** | | | | | | | |
| Software | | | | cryoSPARC-4.5 | | | |
| Particles refinement | 28966 | 39522 | 28752 | 30935 | 29809 | 28225 | 25539 |
| Symmetry | C1 | C1 | C1 | C1 | C1 | C1 | C1 |
| Resolution (Å) | 3.6 | 2.9 | 3.0 | 3.0 | 3.1 | 3.1 | 3.1 |
| Sharpening B-factor (Å$^2$) | 42.5 | 50.0 | 43.6 | 46.0 | 43.7 | 45.4 | 41.6 |
| **Refinement** | | | | | | | |
| **Model composition** | | | | | | | |
| Number of atoms | 32338 | 29703 | 32469 | 32430 | 32402 | 32385 | 32361 |
| Protein residues | 3930 | 3592 | 3943 | 3937 | 3934 | 3932 | 3929 |
| Nucleotides | 53 | 53 | 54 | 54 | 54 | 54 | 54 |
| Ligand | 8 ZN; 1 MG | 1 GTP; 1 MG; 8 ZN | 2 MG; 1 PPI; 8 ZN | 8 ZN; 1 MG; 1 PPI | 1 MG; 1 PPI; 8 ZN | 8 ZN; 1 MG; 1 PPI | 1 MG; 1 PPI; 8 ZN |
| **Bonds RMSD** | | | | | | | |
| Bonds lengths (Å) | 0.012 | 0.0028 | 0.0120 | 0.012 | 0.0118 | 0.012 | 0.0121 |
| Bonds angles (°) | 0.57 | 0.54 | 0.54 | 0.54 | 0.49 | 0.52 | 0.57 |
| **Validation** | | | | | | | |
| MolProbity score | 1.79 | 1.57 | 1.70 | 1.61 | 1.53 | 1.50 | 1.77 |
| Clash score | 9.42 | 6.43 | 8.23 | 7.41 | 6.76 | 7.19 | 9.94 |
| Rotamer outliers | 0.00% | 0.00% | 0.00% | 0.00% | 0.00% | 0.09% | 0.00% |
| C-beta outliers | 0.03% | 0.00% | 1.00% | 0.03% | 1.00% | 0.03% | 1.00% |
| **Ramachandran plot** | | | | | | | |
| Favored (%) | 95.85% | 96.67% | 96.22% | 96.71% | 97.11% | 97.50% | 96.23% |
| Allowed (%) | 4.05% | 3.19% | 3.70% | 3.24% | 2.84% | 2.47% | 3.74% |
| Outlier (%) | 0.10% | 0.14% | 0.08% | 0.05% | 0.05% | 0.03% | 0.03% |

**Table S5 | Cryo-EM data collection and structure determination of Long-RNA RNAPII EC with substrate ATP (Dataset 4)**

| Map | State III (TL-open) | State V (TL-closed) |
|---|---|---|
| **PDB** | 30EP | 30ES |
| **EMDB** | EMD-57673 | EMD-57676 |
| **Data collection** | | |
| Microscope | Titan Krios Gatan | |
| Detector | K3 with GIF Quantum | |
| Cs (mm) | 2.7 | 2.7 |
| Magnification | 105k | 105k |
| Pixel size (Å) | 0.932 | 0.932 |
| Electron dose (e-/Å$^2$) | 50 | 50 |
| Defocus range (μm) | -0.5 to -2.5 | -0.5 to -2.5 |
| Micrograph Number | 2000 | 2000 |
| **Reconstruction** | | |
| Software | cryoSPARC-4.5 | |
| Particles refinement | | |
| Symmetry | | |
| Resolution (Å) | | |
| Sharpening B-factor (Å$^2$) | | |
| **Refinement** | | |
| **Model composition** | | |
| Number of atoms | 30347 | 30516 |
| Protein residues | 3558 | 3570 |
| Nucleotides | 98 | 102 |
| Ligand | 1 ATP; 2 MG; 8 ZN | 1 ATP; 2 MG; 8 ZN |
| **Bonds RMSD** | | |
| Bonds lengths (Å) | 0.0237 | 0.0236 |
| Bonds angles (°) | 0.96 | 0.93 |
| **Validation** | | |
| MolProbity score | 1.69 | 1.60 |
| Clash score | 8.09 | 7.43 |
| Rotamer outliers | 0.13% | 0.16% |
| C-beta outliers | 0.00% | 0.00% |
| **Ramachandran plot** | | |
| Favored (%) | 96.32% | 96.82% |
| Allowed (%) | 3.54% | 2.95% |
| Outlier (%) | 0.14% | 0.23% |

**Table S6 | Summary of RNAPII EC elements with residue ranges and color codes**

| Domain & Motif | Subunits | Residue range | Color code (Default) |
|---|---|---|---|
| Module 1 | Rpb1 (Major), Rpb2, Rpb4–7 | Rpb1: 1–662, 809–1141, 1275–end; Rpb2: 1127–1224; Rpb4–7 (full-length) | Light blue |
| Module 2 | Rpb1, Rpb2 (Major), Rpb3, Rpb9–12 | Rpb1: 663–808, 1142–1274; Rpb2: 1–1126; Rpb3, Rpb9–12 (full-length) | Tan |
| Clamp (include Clamp head and core) | Rpb1 | Rpb1: 1–346, 1396–1436 | Pale goldenrod |
| Lid | Rpb1 | Rpb1: 250–260 | Black |
| Funnel | Rpb1 | Rpb1: 664–808 | Orange or Orange red |
| Funnel tip | Rpb1 | Rpb1: 706–708 | Red |
| Cleft | Rpb1 | Rpb1: 809–871, 1060–1141, 1276–1395 | Yellow |
| Bridge helix (BH) | Rpb1 | Rpb1: 809–848 | Light green |
| Trigger loop (TL) | Rpb1 | Rpb1: 1063–1107 | Purple |
| Protrusion | Rpb2 | Rpb2: 45–218, 406–465 | Goldenrod |
| Lobe | Rpb2 | Rpb2: 219–405 | Sienna |
| Hybrid-binding (HB) | Rpb2 | Rpb2: 751–852, 974–1127 | Steel blue |
| Wall | Rpb2 | Rpb2: 853–973 | Navy |
| Lower jaw | Rpb5 | Rpb5: 1–145 | Slate blue |
| Upper jaw | Rpb1, Rpb9 | Rpb1: 1142–1275; Rpb9: 1-39 | Slate blue |
| Template strand DNA (tsDNA) | NA | NA | Cyan |
| Non-template strand DNA (ntsDNA) | NA | NA | Lime |
| RNA | NA | NA | Hot pink |
| ATP | NA | NA | Gold or hot pink |
| $Mg^{2+}$ ions | NA | NA | Green |

### Extended Data Fig. 1

#### Dataset 1

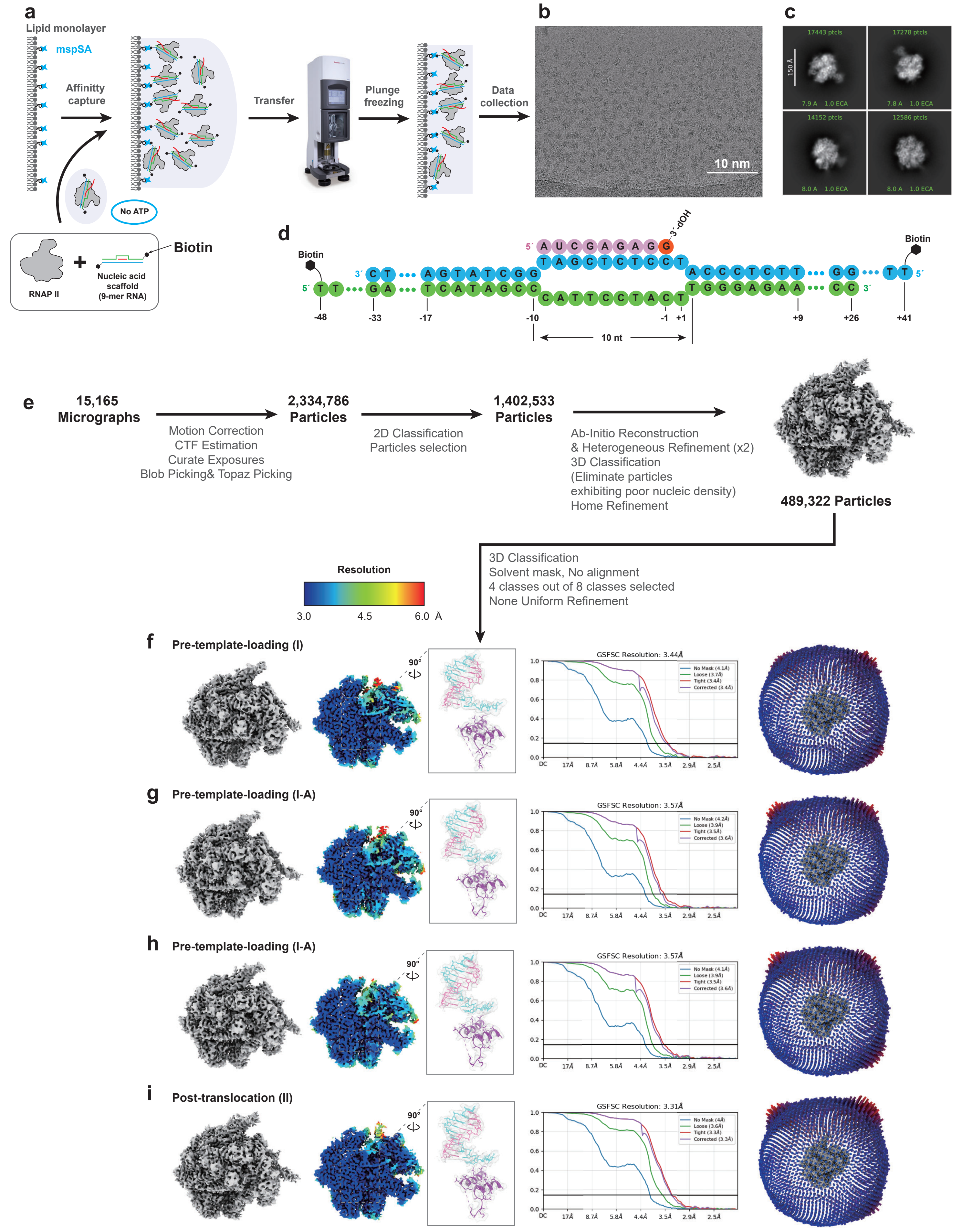

### Extended Data Fig. 2

#### Dataset 2

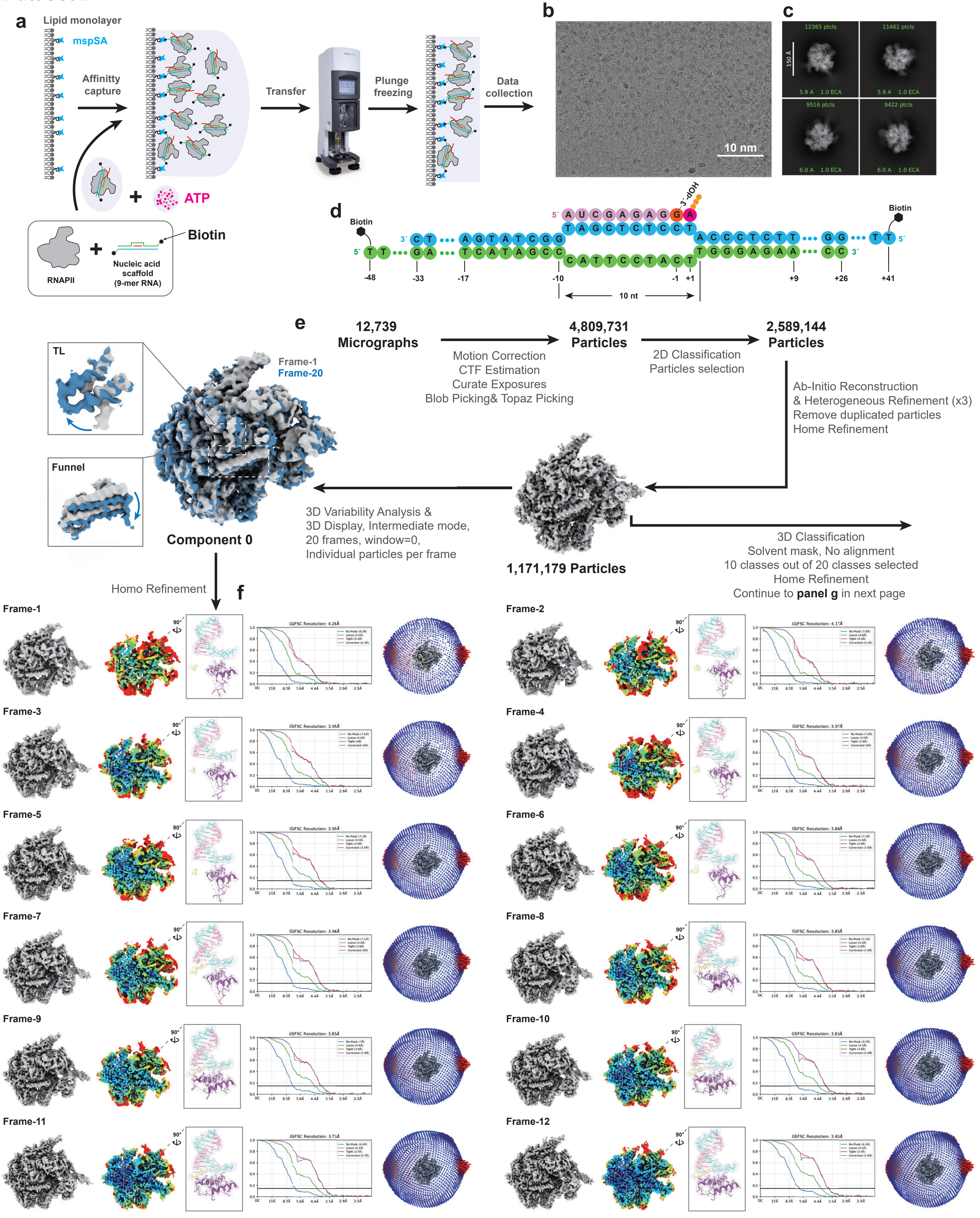

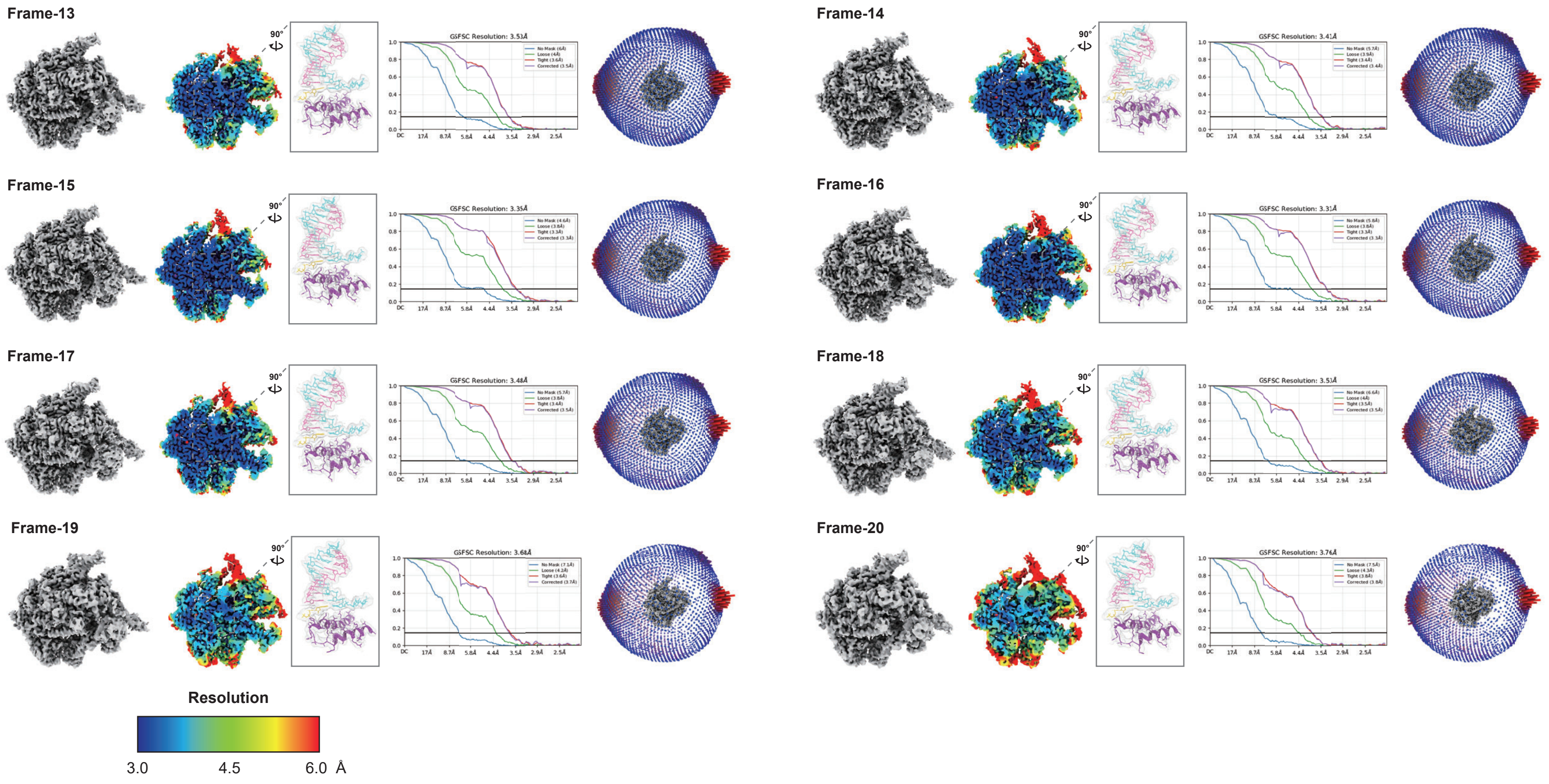


**g**

Continued processing for Extended Data Fig. 2d
3D Classification & Home Refinement

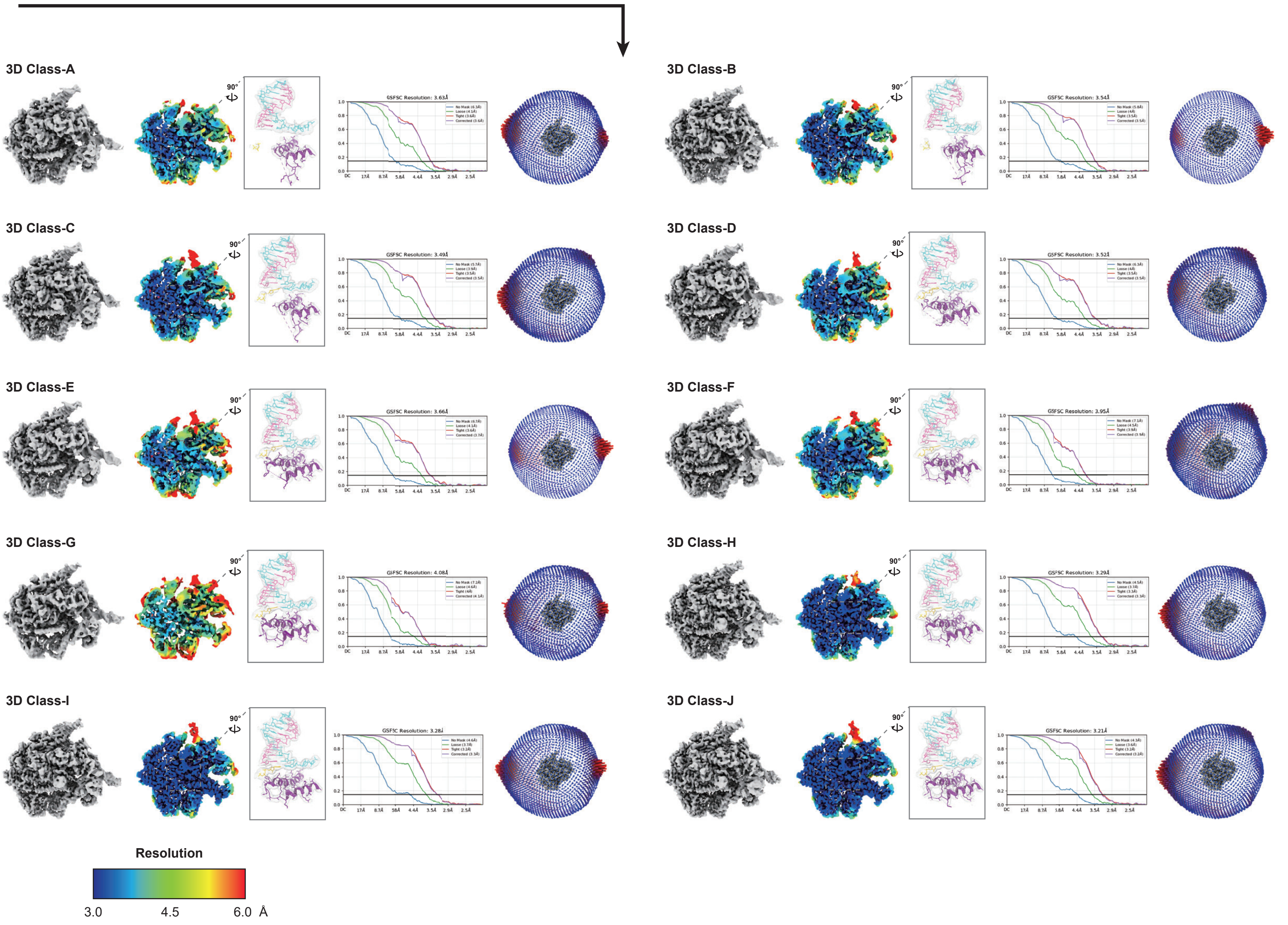

#### Extended Data Fig. 3

#### Dataset 3

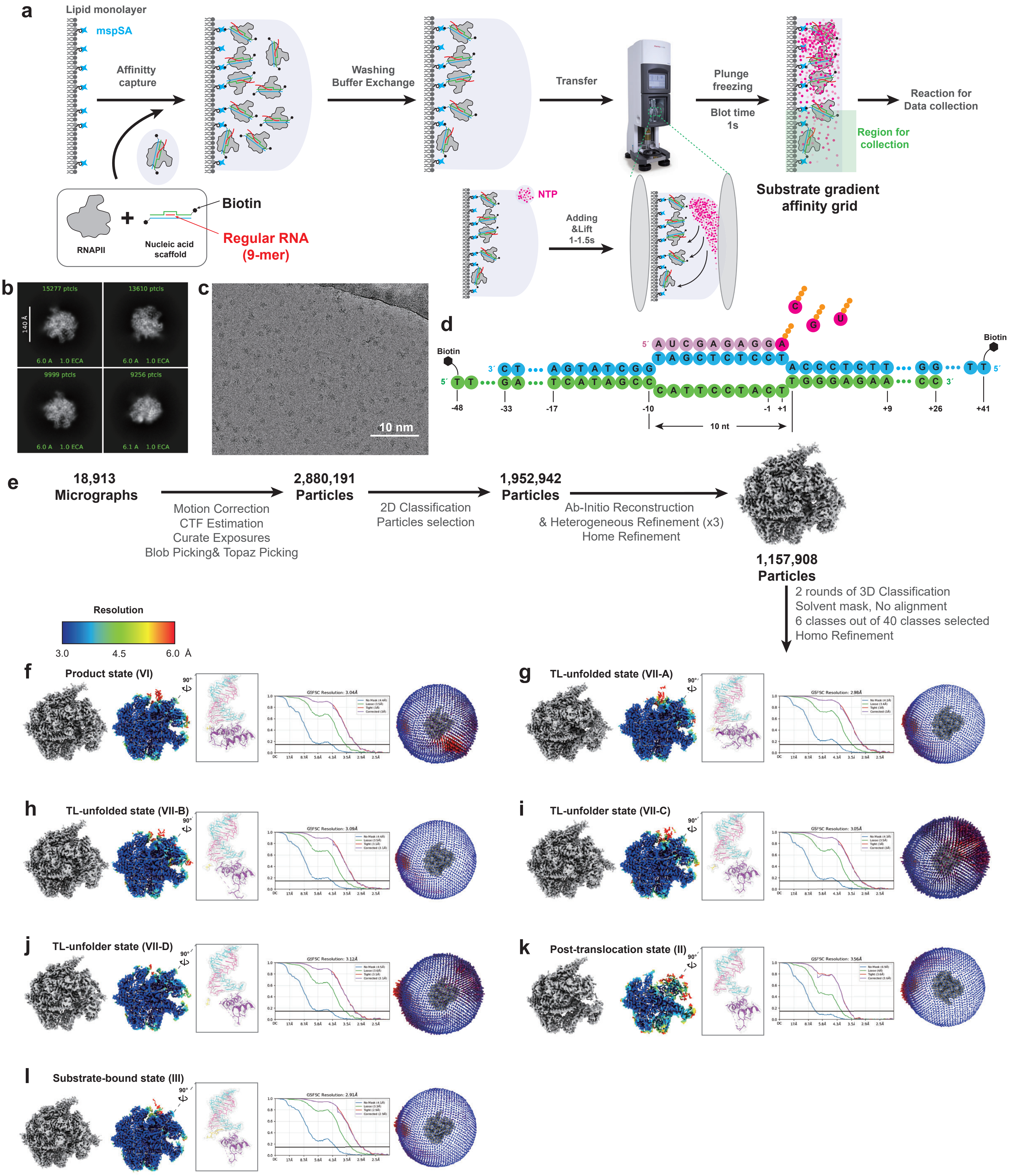

### Extended Data Fig. 4

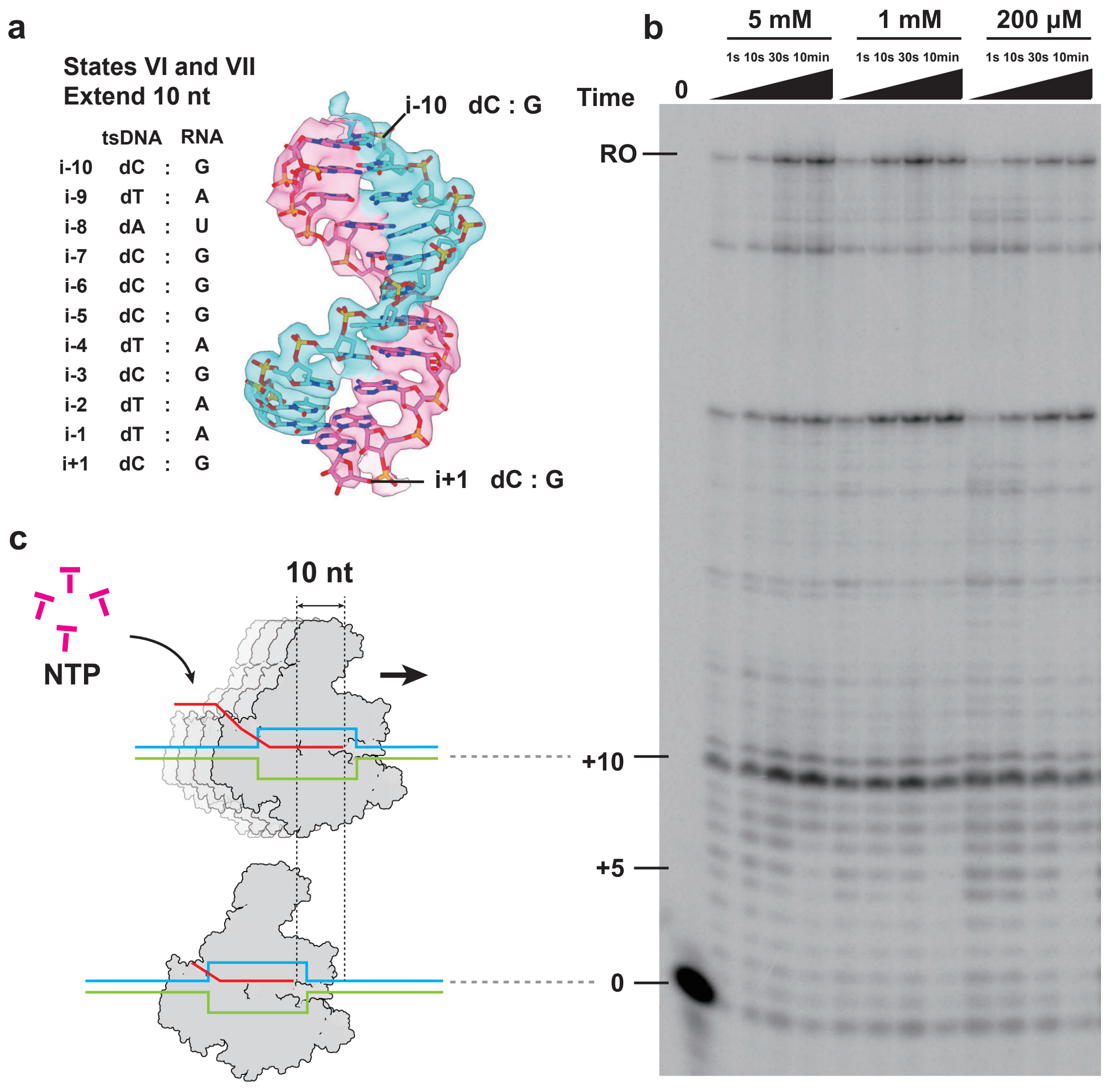

#### Extended Data Fig. 5

#### Dataset 4

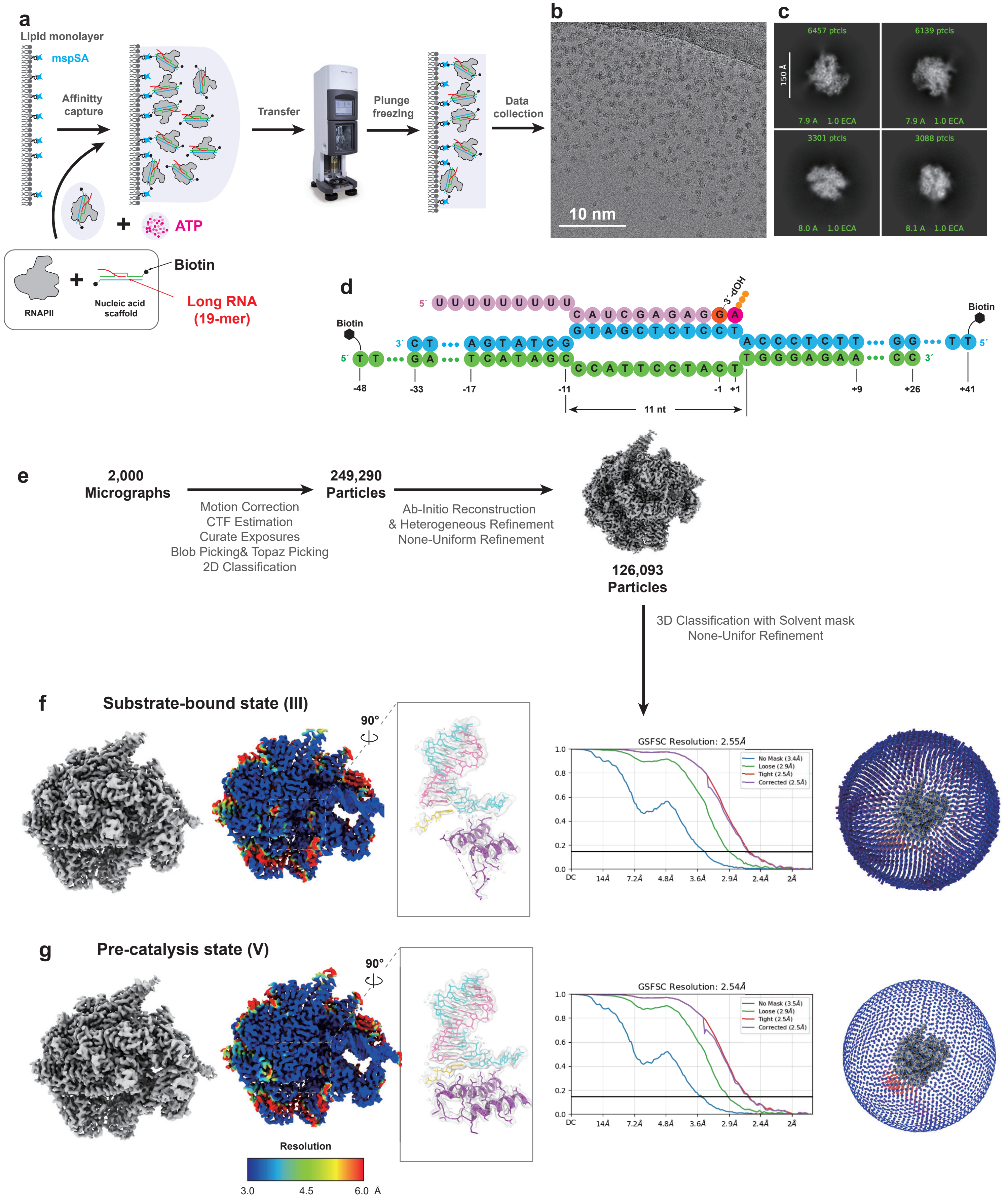

Extended Data Fig. 6

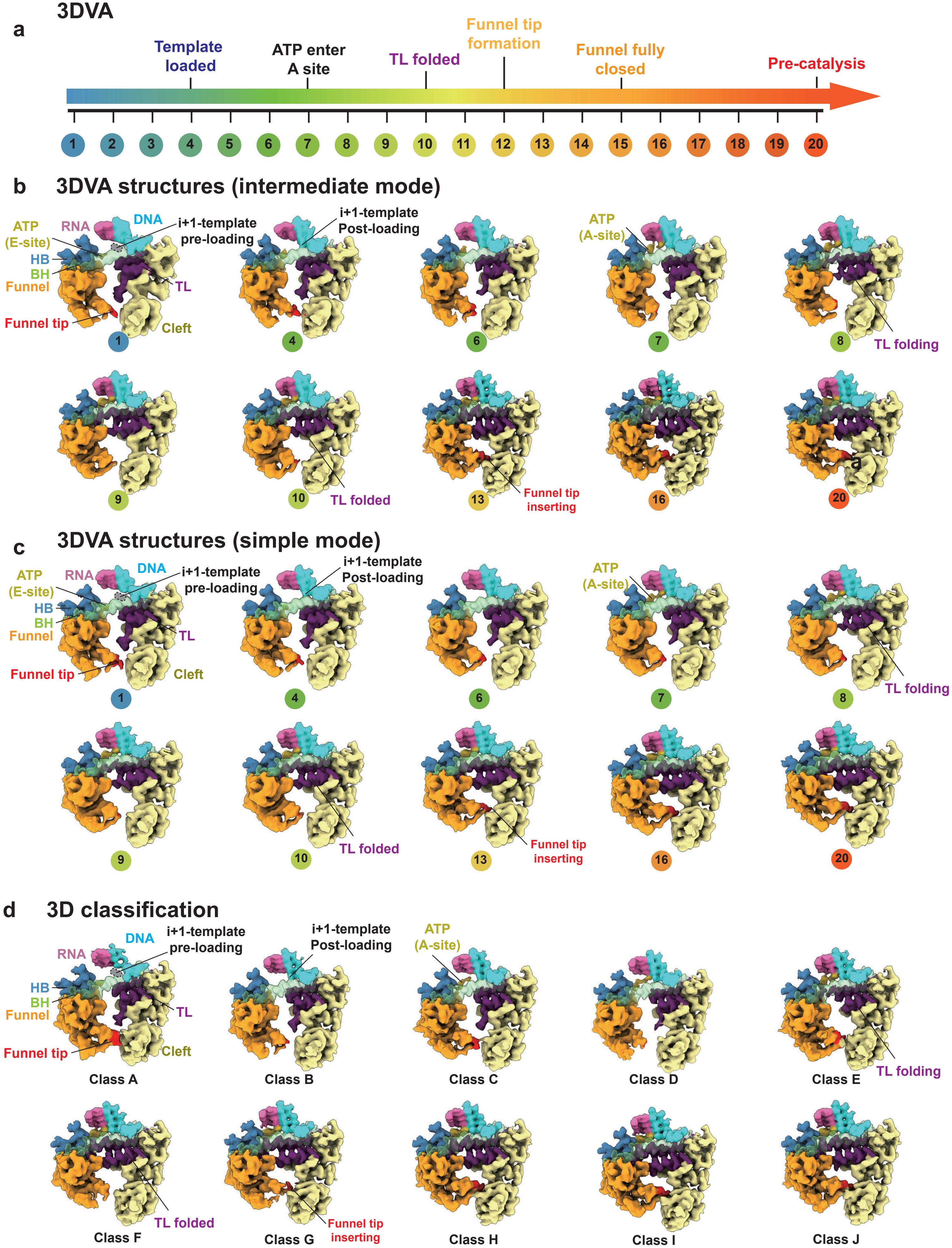

Extended Data Fig. 7

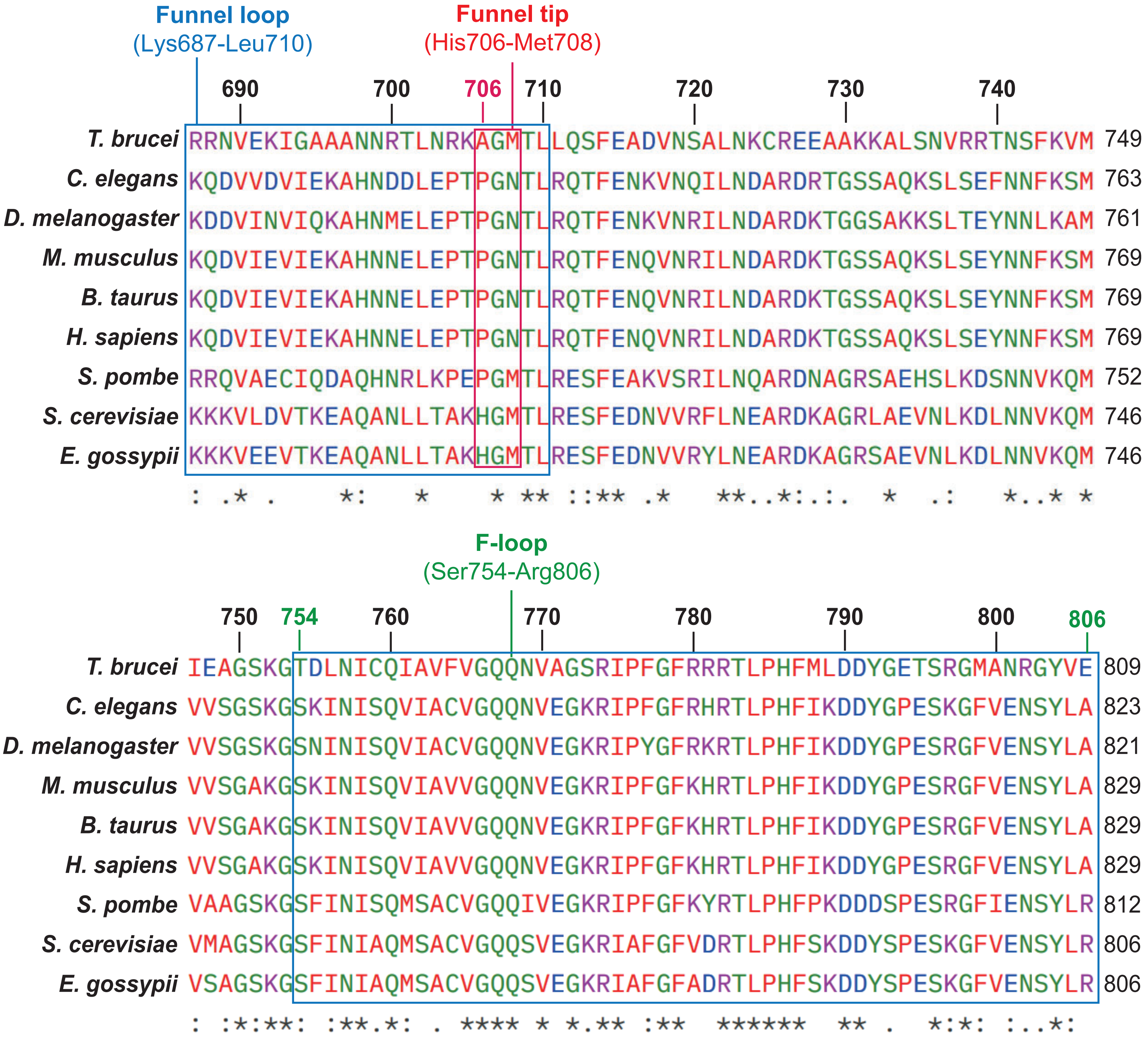

#### Extended Data Fig. 8

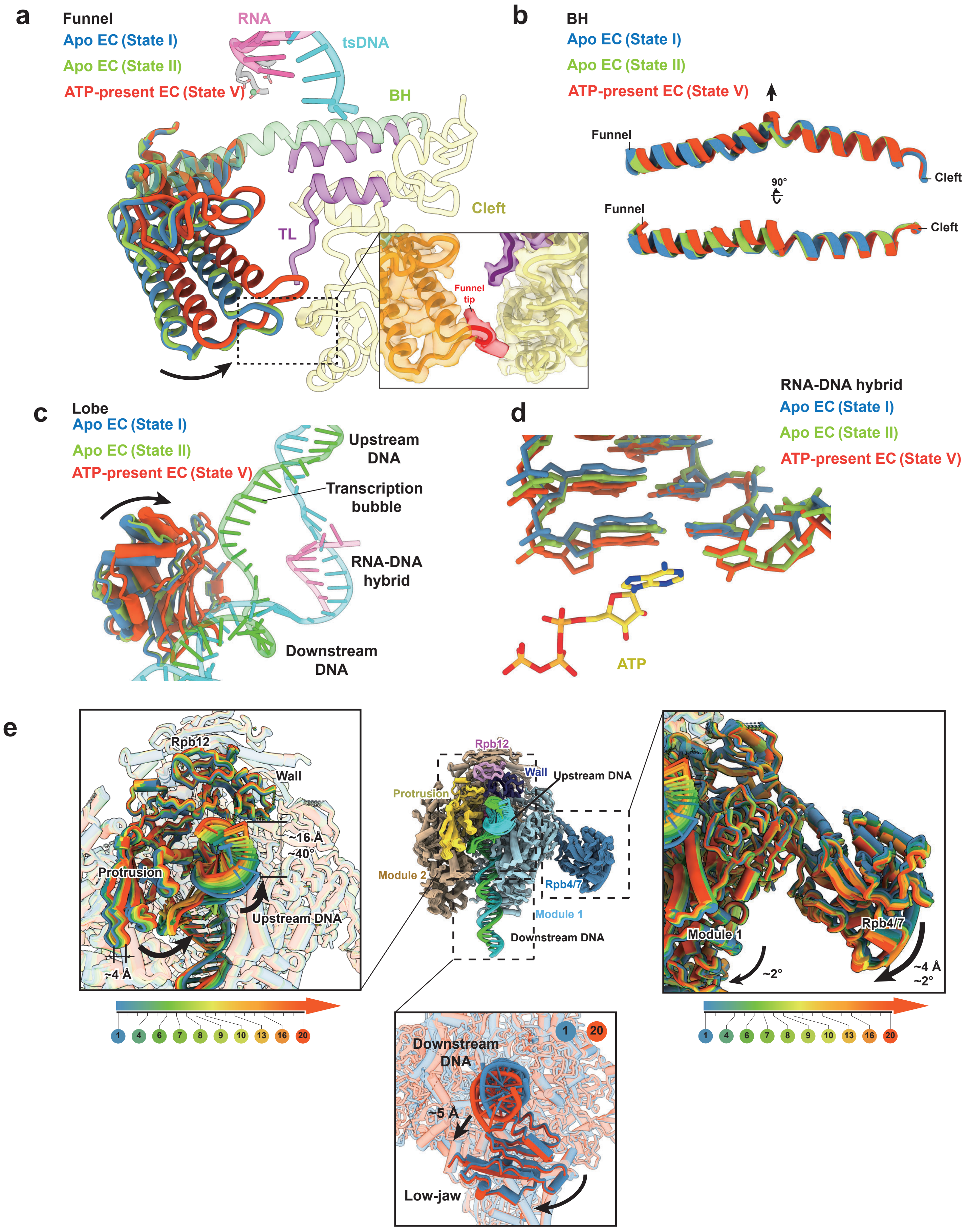

#### Extended Data Fig. 9

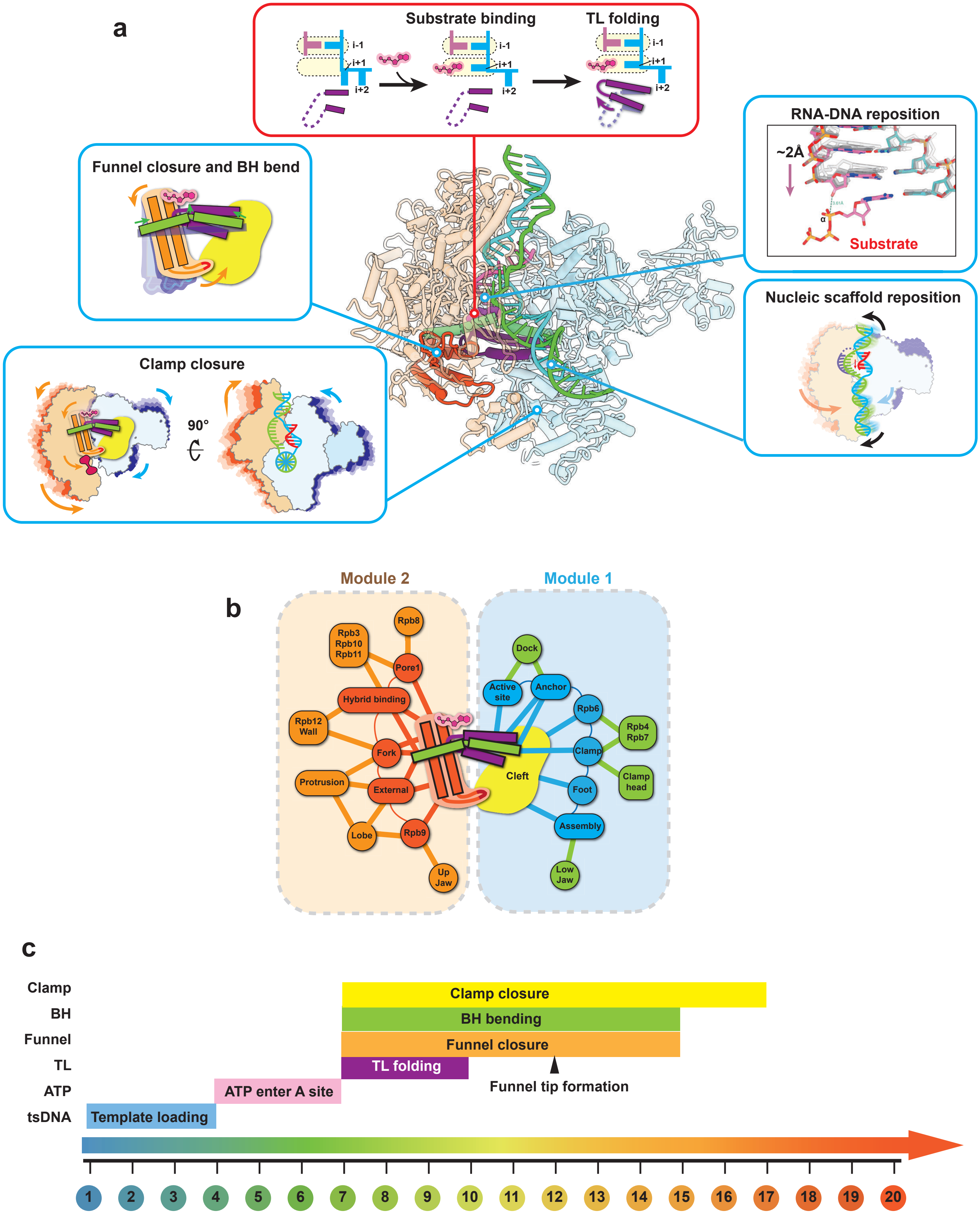

#### Extended Data Fig. 10

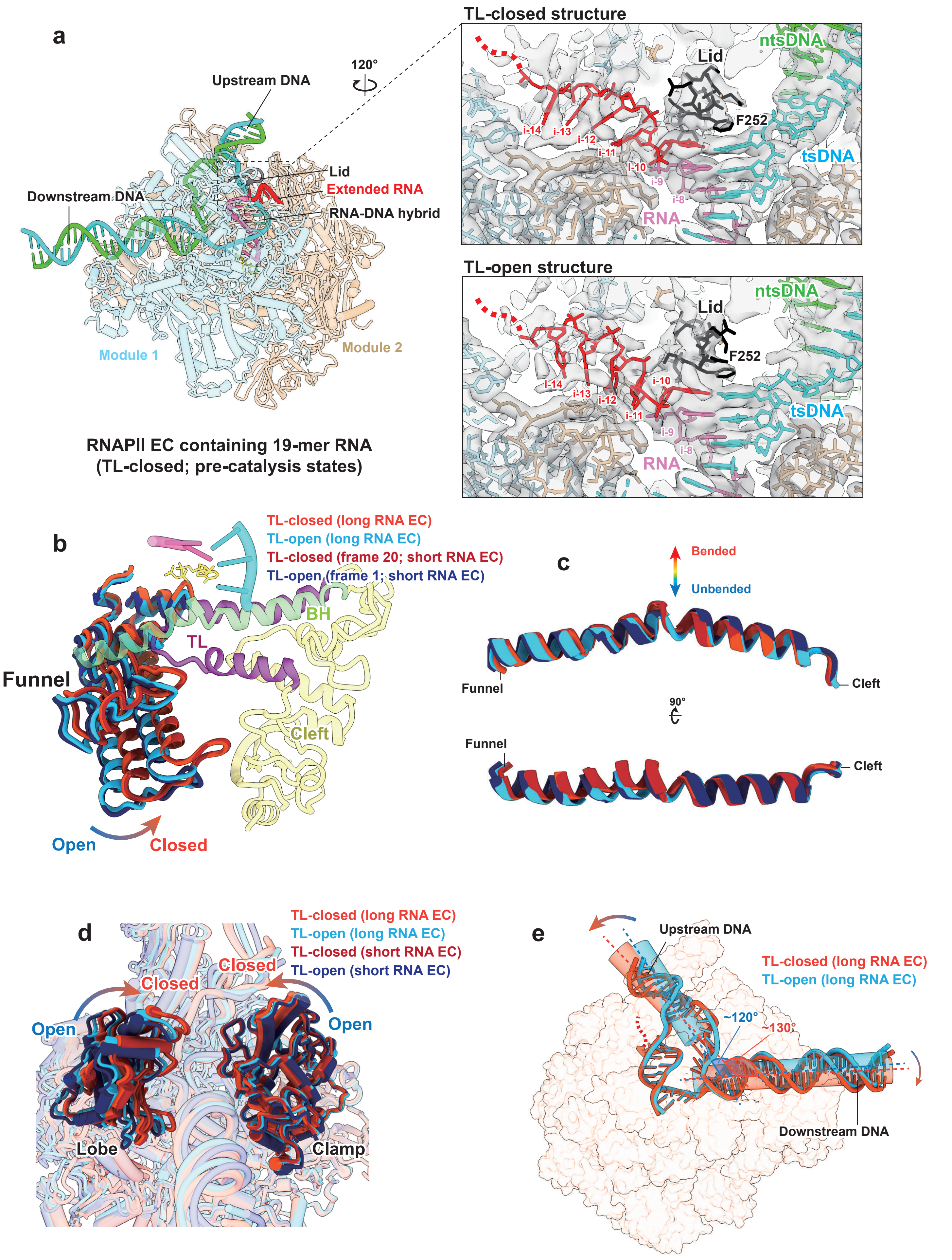

**Extended Data Fig. 11**

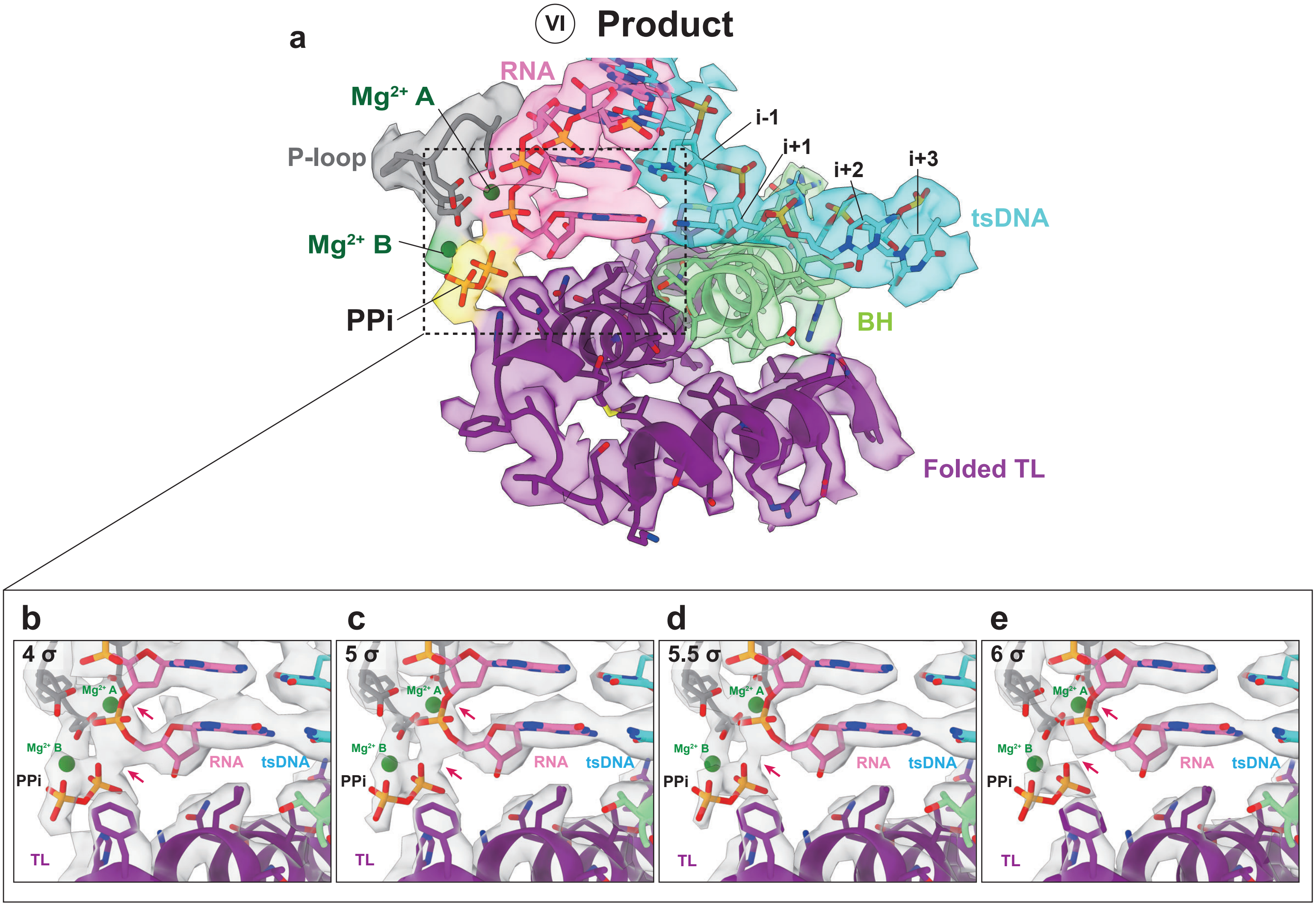

#### Extended Data Fig. 12

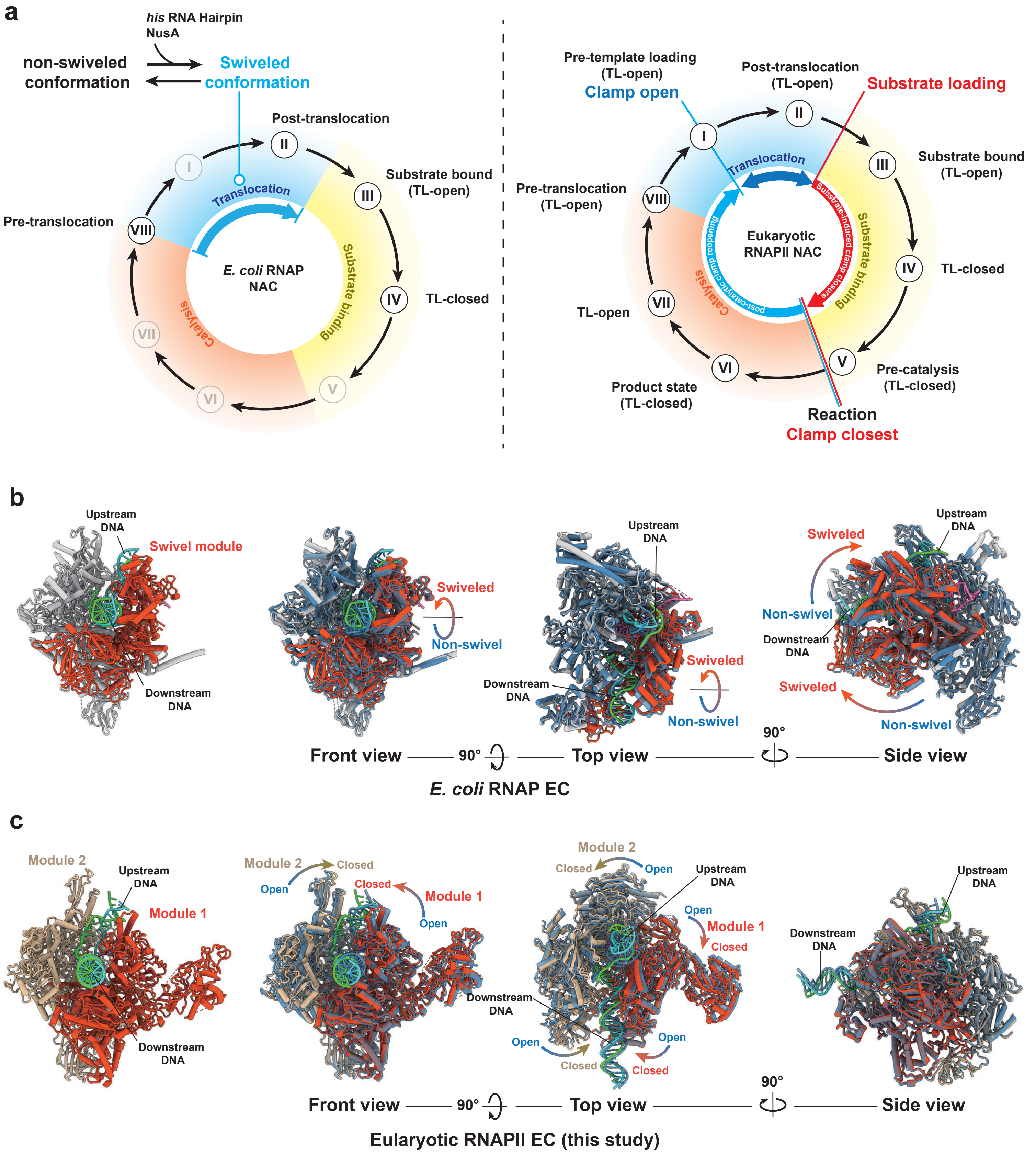

**Extended Data Figure Legends:**

**Extended Data Fig. 1 | Data processing of cryo-EM structures of RNAPII EC without substrate (Dataset 1).**
(a) Schematic of mspSA affinity grid preparation and cryo-EM data collection for Dataset 1. The nucleic acid scaffold was biotinylated to immobilize the RNAPII EC on the grid, thereby reducing particle orientation bias.
(b) Representative raw micrographs of the RNAPII EC.
(c) Gallery of reference-free 2D class averages of the RNAPII EC.
(d) Cartoon scheme for the nucleic acid scaffold used in Dataset 1.
(e) Data processing workflow for Dataset 1.
(f–i) RNAPII EC structures were determined in this dataset. Each structure comprises five representations arranged from left to right: the cryo-EM density map, a slice view colored by local resolution, a close-up of the active center, the Fourier shell correlation (FSC) curve, and the angular distribution plot. In each close-up view, it contains the TL (purple), RNA primer and ATP substrate (hot pink), and template DNA (cyan).

**Extended Data Fig. 2 | Data processing of cryo-EM structures of RNAPII EC with substrate ATP (Dataset 2)**
(a) Schematic of mspSA affinity grid preparation and cryo-EM data collection for Dataset 2. Substrate ATP was included to capture conformational dynamics in the pre-catalytic phase.
(b) Representative raw micrographs of the RNAPII EC with ATP.
(c) Gallery of reference-free 2D class averages of the RNAPII EC containing substrate ATP.
(d) Cartoon scheme for the nucleic acid scaffold used in the cryo-EM dataset of RNAPII EC with substrate ATP.
(e) Data processing workflow for the RNAPII EC substrate binding dataset.
(f–g) RNAPII EC structures were determined from 3D Variability Analysis (f, from Frame 1–20) and 3D classification with a solvent mask (g, from class A–J). Each structure comprises five representations arranged from left to right: the cryo-EM density map, a slice view colored by local resolution, a close-up of the active center, the Fourier shell correlation (FSC) curve, and the angular distribution plot. In each close-up view, it contains the TL (purple), RNA primer and ATP substrate (hot pink), and template DNA (cyan).

**Extended Data Fig. 3 | Data processing of cryo-EM structures of RNAPII EC with substrate NTP (Dataset 3)**
(a) Schematic of substrate-gradient grid (SGG) preparation and cryo-EM data collection for Dataset 3. A regular 9-mer RNA scaffold and substrate NTPs (ATP, UTP, CTP, and GTP) were used to enable RNA extension and capture conformational states in the post-catalytic phase. (See Methods for details of substrate-gradient grid preparation.)
(b) Representative raw micrographs.
(c) Gallery of reference-free 2D class averages.
(d) Cartoon scheme for the nuclear acid scaffold used in Dataset 3.

(e) Data processing workflow.
(f–i) RNAPII EC structures were determined in Dataset 3. Each structure comprises five representations arranged from left to right: the cryo-EM density map, a slice view colored by local resolution, a close-up of the active center, the Fourier shell correlation (FSC) curve, and the angular distribution plot. In each close-up view, it contains the TL (purple), RNA primer and ATP substrate (hot pink), and template DNA (cyan).

**Extended Data Fig. 4 | Transcription assays of RNAPII EC incorporating NTP**
(a) RNAPII EC extends RNA by 10 nucleotides (nt) upon NTP addition, as observed in the post-catalysis cryo-EM dataset. The corresponding RNA-DNA hybrid sequence for states VI and VII is shown on the left. The cryo-EM structure of the RNA-DNA hybrid in RNAPII EC (State VI) is presented on the right, visualized as densities and stick models. Template DNA and RNA are colored cyan and hot pink, respectively.
(b) Transcription assays of RNAPII EC conducted with substrate NTP at three different concentrations: 5 mM, 1 mM, and 200 μM. For each concentration, reactions were quenched at various time points: 5 s, 10 s, 30 s, and 10 min. Markers indicating nucleotide addition are annotated on the side of the gel, and the run-off product is labeled as RO.
(c) Cartoon schematics depicting the elongation of RNAPII EC, illustrating that the elongation complex predominantly pauses after extending by 10 nucleotides.

**Extended Data Fig. 5 | Data processing of cryo-EM structures of long-RNA EC with substrate ATP (Dataset 4)**
(a) Schematic of mspSA affinity grid preparation and cryo-EM data collection for Dataset 4. A long RNA (19-mer) was used to assemble the EC, mimicking a mature EC and enabling cross-validation with Dataset 2.
(b) Representative raw micrographs.
(c) Gallery of reference-free 2D class averages.
(d) Cartoon scheme for the nuclear acid scaffold used in the Dataset 4.
(e) Data processing workflow.
(f–g) Two RNAPII EC structures were determined in Dataset 4. Each structure comprises five representations arranged from left to right: the cryo-EM density map, a slice view colored by local resolution, a close-up of the active center, the Fourier shell correlation (FSC) curve, and the angular distribution plot. In each close-up view, it contains the TL (purple), RNA primer and ATP substrate (hot pink), and template DNA (cyan).

**Extended Data Fig. 6 | Validation of RNAPII EC conformational dynamic cascade in the presence of ATP**
(a) Spectrum illustrating the conformational transitions captured by 3D variability analysis (3DVA) structures from Frame 1 (blue) to Frame 20 (red). Key conformational events associated with specific frames are annotated.
(b–c) Cryo-EM structures of the active center of RNAPII EC, captured at various 3DVA frames derived from two distinct data processing approaches: intermediate mode (b) and simple mode (c).

Structures are displayed as cryo-EM densities and colored according to RNAPII EC components: RNA (hot pink), template DNA (cyan), bridge helix (BH, light green), hybrid binding domain (HB, steel blue), TL (purple), funnel (orange), ATP (gold), and funnel tip (red). Frame numbers corresponding to each structure are indicated below. Key conformational events are annotated alongside the structures for clarity.

(d) Cryo-EM structures generated through 3D classification. These structures illustrate conformational events similar to those captured by 3DVA structures. The 3D classes A-J correspond to respective frame numbers in the 3DVA (b, c).

**Extended Data Fig. 7 | Multi-sequence alignment of funnel tip**

Protein sequences of Rpb1 are aligned by the program of Clustal Omega (1.2.4)[33]. There are nine Rpb1 sequences compared here, including *Saccharomyces cerevisiae*, *Schizosaccharomyces pombe*, *Eremothecium gossypii*, *Trypanosoma brucei*, *Caenorhabditis elegans*, *Drosophila melanogaster*, *Bos taurus*, *Mus musculus*, and *Homo sapiens*. The color codes for amino acid letters are based on their polarity and electrostatics. The funnel tip and loop regions are highlighted with red and blue cycles, respectively.

**Extended Data Fig. 8 | Structural insights of substrate-induced EC tightening**

(a–d) Superposition of funnel domain (a), bridge helix (b), lobe (c), and RNA-DNA hybrid (d) across three structures: the pre-template-loading state (state I, blue) and the post-translocation state (state II, light green) from the apo EC dataset (Dataset 1), and the pre-catalysis state structure (state V, red, Frame 20) from the substrate ATP-bound EC dataset (Dataset 2). The zoom-in panel in panel (a) shows cryo-EM densities of the funnel domain with an emphasis on the funnel tip region highlighted in red. Other RNAPII domains are depicted as transparent ribbons, with domain name annotated in matching colors for clarity.

**(e)** Conformational dynamics of Rpb4/7 stalk, upstream DNA and downstream DNA. The middle panel shows the top view of RNAPII EC, demonstrating conformational dynamics across selected frames from the Dataset 2. Multiple structural frames (frames 1, 4, 6, 7, 8, 9, 10, 13, 16, and 20) are superimposed to illustrate the overall range of motion. Structural elements are color-coded as follows: module 1 (light blue), module 2 (tan), protrusion domain (gold), wall domain (navy), and Rpb12 (plum). Three close-up views of the Rpb4/7 stalk (right) downstream DNA, and upstream DNA (left) correspond to the local region indicated in middle panel. Each close-up view shows the superposition of selected frames as indicated, which are colored cording to the dynamics spectrum from blue (open) to red (closed). They illustrate the gradual and continuous movement of the Rpb4/7 stalk or upstream DNA during the conformational transition. The direction of motion is indicated by black arrows, and the corresponding displacement and rotation are annotated accordingly.

**Extended Data Fig. 9 | Additional analysis of substrate-induced EC tightening**

(a) Schematic representation of coordinated conformational changes in RNAPII during catalysis. A central RNAPII structure is shown with key structural elements highlighted, including the substrate (magenta), TL (purple), BH (light green), funnel (orange red), module 1 (light blue) and module 2 (tan). Surrounding cartoon diagrams illustrate the conformational changes of individual elements. Each schematic is connected to its corresponding structural location by lines, emphasizing spatial mapping rather than causal relationships. Previously reported conformational changes are indicated by red boxes, whereas newly identified changes in this study are highlighted in blue boxes.

(b) Interaction network underlying RNAPII conformational dynamics. The network is centered on the catalytic region, including the substrate, TL, funnel, BH and Cleft. RNAPII is divided into two modules, module 1 and 2. First-layer interactions are highlighted within module 2 (dark orange) and module 1 (sky blue), representing elements directly connected to the core region. Peripheral interactions are further indicated in module 2 (light orange) and module 1 (light green), illustrating an extended interaction network.

(c) Conformational dynamics spectrum derived from the Dataset 2. A series of 20 structural frames (states I–V and intermediates) are arranged to represent the continuous transition from an open to a tightened RNAPII conformation. Key conformational dynamic events are mapped onto this spectrum. The intervals highlighted for each structural element correspond to the frames in which the most pronounced conformational changes occur. These assignments are based on comparative analyses of local structural deviations between frames. Importantly, the indicated intervals represent regions of dominant structural rearrangement rather than the full extent of conformational variation. Structural elements may continue to undergo more subtle adjustments outside these intervals, including side-chain rearrangements or local density differences, which are not explicitly highlighted in this representation.

**Extended Data Fig. 10 | Cryo-EM structures of RNAPII EC containing long RNA (19-mer)**

(a) Overall structure of the RNAPII elongation complex (EC) assembled with an extended 19-mer RNA in the TL-closed state, representing a more complete elongation state compared to short-RNA complexes. The left panel shows the global architecture of the EC, with module 1 colored light blue and module 2 in tan. The nucleic acid scaffold includes tsDNA (cyan), ntsDNA (lime), and RNA (hot pink), with the extended upstream RNA highlighted in red. The right panel shows zoom-in views of the upstream RNA junction in two conformational states: TL-closed (upper) and TL-open (lower). Cryo-EM densities and corresponding atomic models illustrate RNA extension and strand separation from the tsDNA. The lid domain is colored black, with residue F252 highlighted.

(b–d) Structural comparisons of key dynamic elements across four RNAPII EC structures. The funnel domain (b), bridge helix (c), and lobe/clamp domain (d) are compared between long-RNA dataset 4 (TL-closed, orange red; TL-open, sky blue) and short-RNA dataset 2 (frame 20, firebrick; frame 1, navy). A continuous color gradient from blue (open) to red (closed) illustrates domain movements.

(e) Nucleic acid scaffold rearrangements accompanying EC tightening in the long-RNA dataset. Comparison of TL-open (sky blue) and TL-closed (orange red) states reveals coordinated clamp closure and global nucleic acid repositioning. Arrows indicate directional movements of upstream and downstream DNA. The axes of upstream and downstream DNA are represented by shaded cylinders (blue and red, respectively), highlighting changes in DNA trajectory.

**Extended Data Fig. 11 | Validation of the product state by cryo-EM density maps.**
(a) Cryo-EM structure of the RNA polymerase II elongation complex (EC) captured in the product state (VI). The density map is shown at a contour level of 0.132 (3-σ). Color coding of RNAPII elements: trigger loop (TL, purple), bridge helix (BH, light green), template DNA (tsDNA, cyan), and RNA primer (hot pink).
(b-e) Cryo-EM densities highlighting the newly formed phosphodiester bond and the released pyrophosphate (PPi), displayed at different contour levels: 0.176 (4-σ, panel b), 0.220 (5-σ, panel (c)), 0.242 (5.5-σ, panel (d)), and 0.264 (6-σ, panel (e)). Red arrows indicate the positions of bond formation and cleavage.

**Extended Data Fig. 12 | Comparison between bacterial RNAP swiveling and eukaryotic RNAPII tightening**
(a) Schematic representation of the nucleotide addition cycle (NAC) in bacterial RNA polymerase (left) and eukaryotic RNA polymerase II (right), with conformational transitions mapped onto distinct stages of the cycle. The NAC is divided into three major phases: translocation (blue), substrate binding (yellow), and catalysis (orange). Roman numerals indicate structurally defined intermediate states. Eight states are resolved for RNAPII, whereas only a subset of states has been structurally characterized in bacterial RNAP; unresolved states are depicted in semi-transparent shading. In bacterial RNAP (left), swiveling is coordinated with the translocation phase of the NAC. This conformational equilibrium can be modulated by transcription factors (e.g., NusA2) or pausing signals such as his RNA hairpins1, which bias the enzyme between non-swiveled and swiveled states. In contrast, eukaryotic RNAPII (right) exhibits a distinct mode. Substrate binding promotes clamp closure (red arrow), progressing toward a maximally closed state near the catalytic step. Following catalysis, the clamp reopens (blue arrow), returning toward a more open conformation. The timing of substrate loading, as well as the most open and most closed clamp conformations, are indicated along the cycle.
(b–c) Structural comparison of bacterial RNAP swiveling and eukaryotic RNAPII clamp dynamics. (b) Bacterial RNAP swiveling. The leftmost panel shows the cryo-EM structure of *E. coli* RNAP in the swiveled state (PDB: 6ASX). The swiveling module (orange red) partially overlaps with module 1 in eukaryotic RNAPII. Three orthogonal views (front, top, and side) depict the reversible transition between the non-swiveled (PDB: 6BJS) and swiveled states, with arrows indicating the direction of motion. (c) Eukaryotic RNAPII clamp closure. By contrast, the leftmost panel shows the cryo-EM structure of the RNAPII elongation complex (EC) in the pre-catalytic state (state V) determined in this study. Two major structural modules are highlighted: module 1 (orange red) and module 2 (light blue). Three orthogonal views (front, top, and side) illustrate the coordinated, progressive movement of these modules during clamp dynamics, with gradient arrows indicating the transition from the open (state I; pre-template loading) to the closed (state V; pre-catalysis) conformation.
